# Dietary Bioactive Compounds Trigger Distinct Epigenetic and Metabolic Reprogramming in *Lactobacillus acidophilus*

**DOI:** 10.1101/2024.08.18.608491

**Authors:** Yanzhuo Kong, Damola O. Adejoro, Philip A. Wescombe, Christopher Winefield, Stephen L.W. On, Nadia L. Mitchell, Arvind Subbaraj, Andrew Saunders, Evelyne Maes, Venkata Chelikani

**Affiliations:** Faculty of Agriculture and Life Sciences, Lincoln University, Lincoln 7647, New Zealand; Yili Innovation Centre Oceania, Lincoln University, Lincoln 7647, New Zealand; National Center of Technology Innovation for Dairy, Hohhot, Inner Mongolia, China; Proteins & Metabolites Team, Beyond Food Innovation Centre of Excellence, AgResearch Ltd, Lincoln, New Zealand; Westland Milk Products, PO Box 138, Rolleston 7614, Canterbury, New Zealand

**Keywords:** N4-Methylcytosine, Epigenomics, Probiotics, Dietary bioactive and Nutrifermentics

## Abstract

*Lactobacillus acidophilus* ATCC 4356 (LA), a key probiotic in the human gut microbiota, offers several health benefits. While dietary bioactive compounds are known to influence gut microbiota, their specific mechanisms remain unclear. This study investigated how certain dietary bioactive compounds impact LA gene expression and metabolism. Results showed each compound produces unique transcriptional, metabolic, proteomic, and epigenetic profiles in LA. Notably, dietary compounds altered the epigenetic landscape through N4-methylcytosine (4mC) modification, a relatively underexplored form of methyl modification that may play a role in regulating gene transcription.

For instance, genistein treatment up-regulated 76 genes and the down-regulated 130 genes in LA. A gene involved in mucus-binding proteins, crucial for bacterial adhesion, was up-regulated 38-fold, likely due to 4mC modifications. Additionally, the gene coding for the melibiose operon regulatory protein increased 78-fold, enhancing melibiose (a prebiotic) production with genistein, but only 1.1-fold with sodium butyrate. This study highlights the potential of dietary compounds for microbial metabolic engineering, offering a non-GMO method for modulating bacterial performance and other biotechnology applications.

**Graphical Abstract:** Dietary compounds are able to modify the epigenetic landscape of *Lactobacillus acidophilus* ATCC 4356, resulting in significant transcriptomic, metabolic, and physiological changes. This approach differs from conventional genetic modification techniques for manipulating microbial strains.

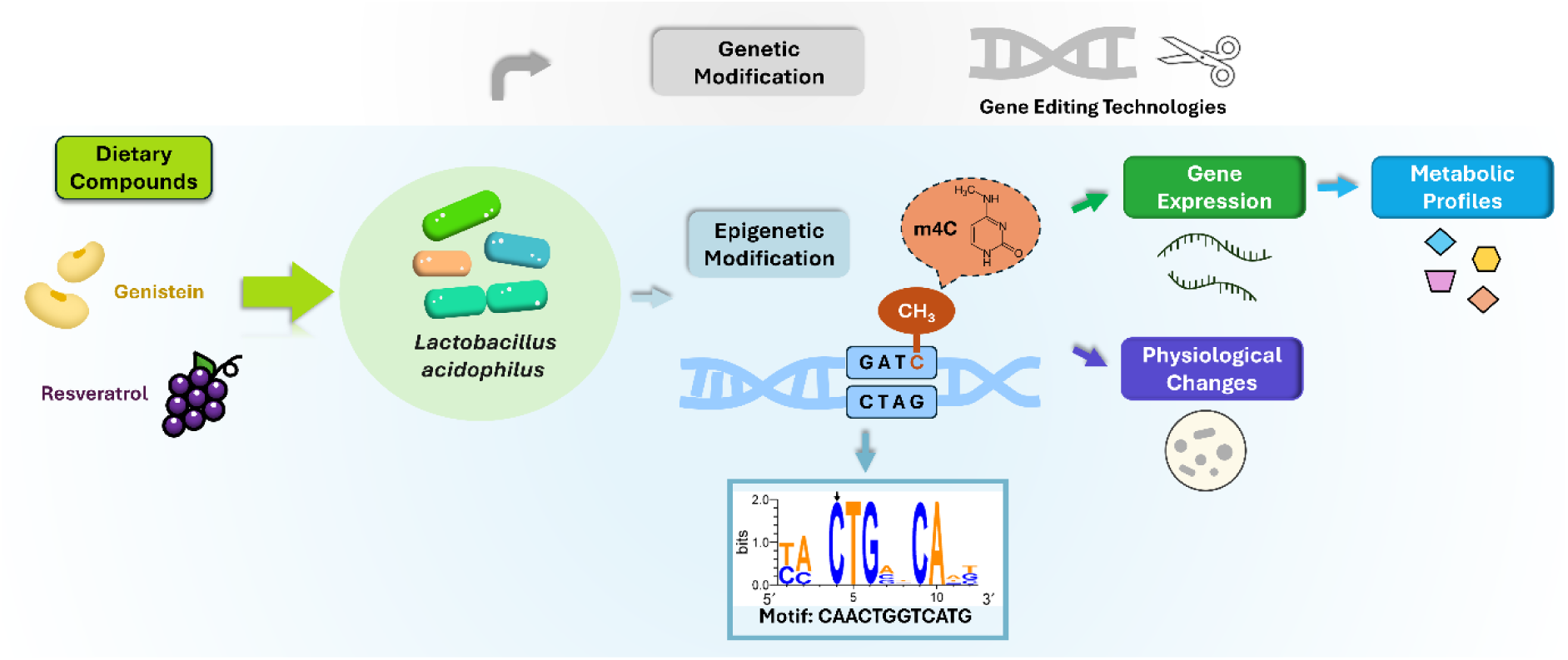

## Introduction

The human gut microbiota plays a vital role in maintaining health and supporting various physiological processes^1^. The *Lactobacillaceae* are important inhabitants of the human gut, and some species are considered to have probiotic properties, offering numerous benefits for health and well-being. Among these, *Lactobacillus acidophilus* ATCC 4356 (LA) is a key probiotic species known for its significant contributions to gut health and is one of the most widely recognized and commercially distributed probiotic cultures^2–4^. Dietary compounds have been shown to affect both the composition and function of the gut microbiota^5–7^. Certain dietary polyphenols and bioactive food compounds like genistein, tea catechin, resveratrol, butyrate, sulforaphane, diallyl sulfide, and curcumin have been associated with inducing epigenetic mechanisms and can inhibit or activate enzymes that catalyze DNA methylation or histone modifications or alter substrate availability, affecting gene expression and physiology^8^. DNA methylation is an epigenetic modification found in both prokaryotic and eukaryotic genomes. In bacteria, modifications to adenine or cytosine, resulting in N6-methyladenine (6mA) or N4-methylcytosine (4mC), respectively, are considered the most significant^9^. Notably, 6mA plays a significant role in regulating gene expression across various processes^10^. Most bacterial DNA methyltransferases (MTases) are components of the restriction-modification (R-M) system. These MTases protect chromosomal DNA from the activity of the corresponding restriction enzyme, which targets and destroys foreign DNA, such as that from bacteriophages^11,12^. However, R-M systems have roles beyond cellular defense; they are also known to influence epigenetics and transcriptome profiles^13^. The occurrence of 4mC is rare in eukaryotes but common in bacteria and archaea^14^. Many bacterial genomes display a 4mC modification, whose physiological significance is not yet clear. There are only a few studies that have investigated the functional role of 4mC modifications, including the role of 4mC modifications in the Type 1 R-M system^15^, secondary metabolism regulation in *Streptomyces roseosporus* L30^14^, and global epigenetic regulation in *Helicobacter pylori*^16^. Alterations in the methylome can result in changes in the transcriptome, which subsequently usually impact phenotypes. These methylome changes offer potential targets for both natural and artificial selection^17^.

Despite extensive research on the roles of diet and the gut microbiota on human health and growing evidence that dietary compounds alter the gut microbiota, there are no comprehensive studies on how dietary compounds influence individual strains. Investigating how these dietary compounds affect microbial transcription and metabolism is essential to uncovering the mechanisms that regulate these metabolic processes. Such knowledge could enable GMO-free metabolic engineering of bacteria for biotechnology applications such as natural product biosynthesis. Current gene editing methods, such as suicide plasmid and Clustered regularly interspaced palindromic repeats (CRISPR)-Cas technologies, involve challenges such as complex laborious, multi-step methods, risk of CRISPR nuclease-associated cytotoxicity, and the outcome is always a genetically modified organism (GMO)^18–23^. Our approach offers potential advantages to current genome editing methods and may also enable improved efficacy of probiotics through epigenomic enhancement, ultimately manipulating gut microbiota functions to enhance human health.

In this present study, we used a multi-omics approach to analyze the transcriptome, metabolome, proteome, and epigenome of LA treated with non-inhibitory concentrations of the bioactive dietary compounds such as genistein and resveratrol. Our findings indicate that each dietary compound exerts unique transcriptional and metabolic effects, and the transcription is potentially regulated by 4mC modifications.

## Results

### Dietary bioactive compounds have unique transcriptomic impacts on *Lactobacillus acidophilus*

A total of 385 genes showed differential expression in the treated strains compared with the untreated wild type (Figure 1). Genistein treatment resulted in the most significantly different changes, with 76 up-regulated genes and 130 down-regulated genes, followed by resveratrol with 27 up-regulated and 68 down-regulated genes (Figure 1b). L-methionine and sodium butyrate had minimal impact, with only 6/18 and 4/3 genes being up-/down-regulated, respectively (Figure 1b).

**Figure 1.**
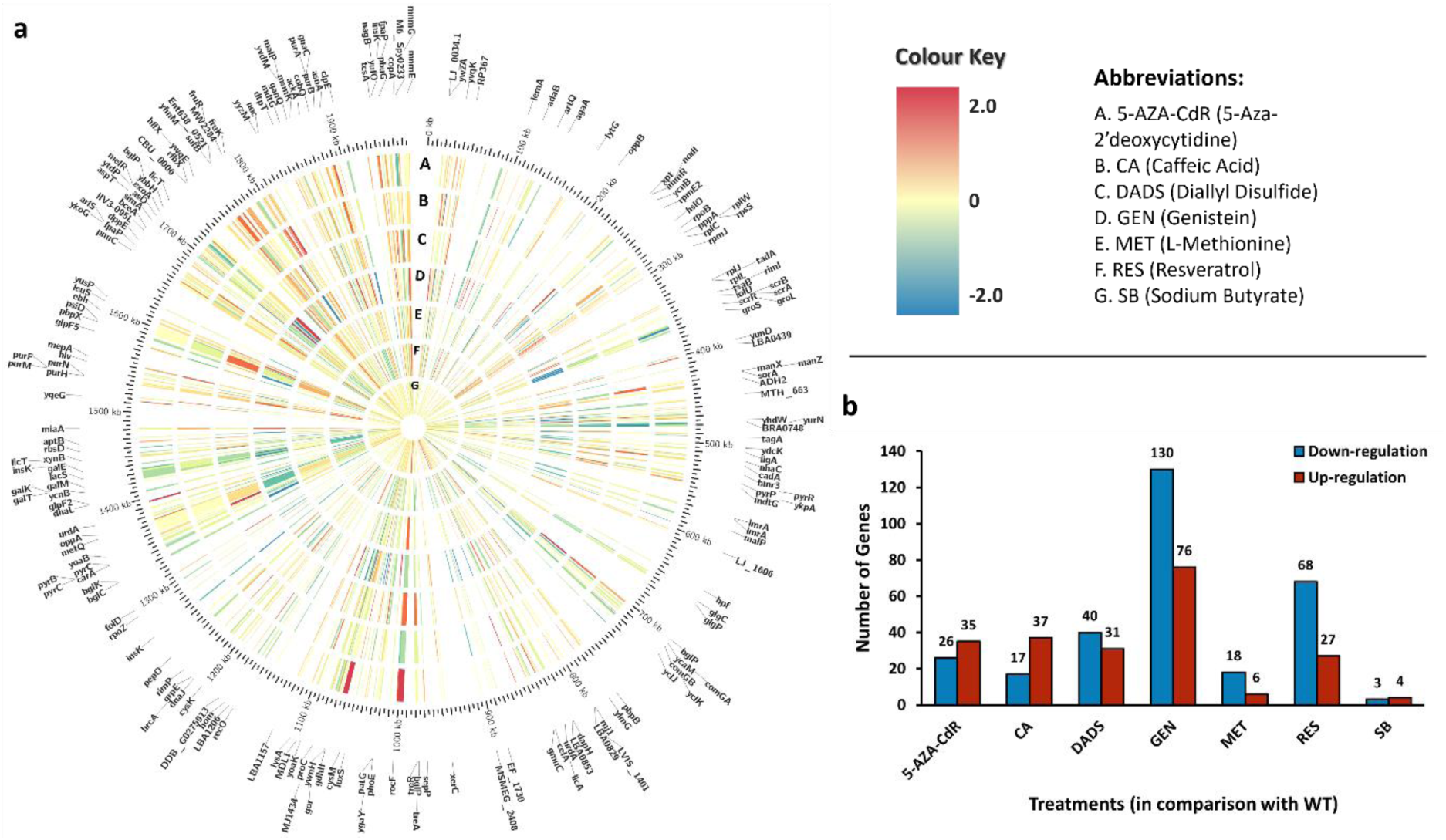
**a.** Circular representation of the *Lactobacillus acidophilus* ATCC 4356 (LA). genome displaying differentially expressed genes (DEGs) as defined by a log_2_ fold change (>1.5 or <-1.5) in expression values determined by RNA sequencing, compared with gene expression levels in the untreated strain. The outer ring lists genes that were differentially expressed when treated with various dietary bioactive compounds. The second ring represents the chromosome and base pair locations (kb). Rings A to G are heatmaps visualizing gene expression changes under the treatment with seven dietary bioactive compounds, listed on the top right. Genome mapping and DEG visualization was done using Circos (version 0.69-8); **b.** The bar graph illustrates the total number of significantly up-regulated and down-regulated genes in LA treated with dietary bioactive compounds compared with the untreated wild type LA. **Abbreviations:** WT: untreated wild type LA; 5-AZA-CdR: 5-Aza-2’deoxycytidine; CA: Caffeic Acid; DADS: Diallyl Disulfide; GEN: Genistein; MET: L-Methionine; RES: Resveratrol; SB: Sodium Butyrate.

Further analysis of the alternatively regulated genes and their likely impact on various metabolic processes was carried out (Extended Data Fig. 1). Each treatment compound elicited a distinct and unique response in LA (Figure 1), with genistein noticeably indicating a predominant shift in carbohydrate metabolism and sugar utilization (Extended Data Fig. 1).

Several genes, including *lysA*, *melR*, *ytdP*, *licT*(2), *ywzA*, *malP*(1), *asnA*, *nagB*, and *mnmG* were significantly up-regulated in genistein-treated strains compared with the untreated wild type, with log_2_ fold changes above 1.5. Notably, *melR*, *ytdP*, and *licT*(2) showed log_2_ fold change values of 8.86, 5.62, and 3.45, respectively. Although the specific roles of *ywzA* and *ytdP* in LA remain poorly understood, many of these genes are known to participate in various metabolic and biosynthetic processes within bacterial cells. For example, *melR* is responsible for the utilization and metabolism of the disaccharide sugar melibiose, similar to *licT*(2), which contributes to carbohydrate metabolism, and *malP*(1) participates in maltose utilization^24–26^. The up-regulation of these genes may reflect a shift in carbohydrate metabolism. In addition, *nagB* has been reported to participate in N-acetylglucosamine (GlcNAc) metabolism, suggesting its involvement in the utilization of this sugar derivative^27,28^. The genes *lysA* and *asnA* are associated with the biosynthesis of the essential amino acid lysine and non-essential amino acid asparagine in LA^29,30^. These observations indicate that the bacterium may respond to dietary compounds by altering its metabolic processes. Furthermore, *mnmG*, involved in tRNA modification in LA, plays a vital role in maintaining the accuracy of codon-anticodon interactions during translation, ensuring proper protein synthesis^31,32^. Overall, these genes represent a wide range of functional categories, highlighting the diversity of biological processes influenced by dietary bioactive compounds in LA.

Conversely, genistein treatment also down-regulated several genes, with many exhibiting log_2_ fold changes below -1.5. This included genes associated with galactose metabolism (*galM* and *galT*)^33^, lactose and peptidoglycan metabolism (*lacZ* and *murQ* respectively)^34,35^, fructose metabolism (*fruA*, *fruK*, and *fruR*)^36,37^ and likely associated with sorbitol metabolism (*scrR* and *scrA*)^38^. The decreased activity of these genes might affect sugar utilization and carbohydrate metabolism in LA. Furthermore, down-regulation of genes like *proC*, which contributes to proline biosynthesis^39^, and those involved in stress responses and cellular maintenance (*clpE*, *groS*, *groL*, and *tagA*), suggests genistein’s impact extends to cellular adaptation and maintenance^40–42^. The precise regulatory mechanisms and downstream effects of these changes require further investigation, but these findings underscore how dietary compounds shape the behavior of LA.

In resveratrol-treated strains, *ywzA* and *mnmG* were significantly up-regulated, like that observed with genistein treatment. The *mnmG* gene is involved in tRNA modification and translation accuracy^31,32^, hence, its up-regulation indicates potential enhancement of translational fidelity and improved synthesis of proteins within LA. However, *galM*, *comGA*, and *licA* were significantly down-regulated. The down-regulation of *galM* and *licA* could potentially affect galactose and peptidoglycan metabolism, altering sugar utilization and affecting bacterial cell wall integrity^33,43^. Down-regulation of *comGA* might indicate a reduced capacity for genetic competence within LA, potentially limiting its ability to incorporate new genetic material and adaptability to novel challenges^44^.

The positive control, 5-aza-2’-deoxycytidine, significantly up-regulated *guaC* and down-regulated *copA* and *scrA*. Up-regulation of *guaC* suggests increased guanine nucleotide biosynthesis, essential for DNA and RNA synthesis, thereby supporting cell growth and division^45^. Conversely, down-regulation of *copA* and *scrA* may signify alterations in the metabolism or transport of specific compounds like excess copper ions and sorbitol, with broader implications for the overall metabolism of LA^38,46^. Caffeic acid led to the significant down-regulation of *yurN* and *yesQ*, with unclear effects due to the limited information available about these genes, highlighting the need for further research to uncover their functions and associated biological pathways. Diallyl disulfide treatment up-regulated *yoaB*, *fpaP*(2), *immR*, and *bglP*(2), and down-regulated *insK1*. Up-regulation of *bglP*(2), the beta-glucoside gene, might indicate an increased capacity of LA to transport and utilize sugars, especially beta-glucoside, as carbon sources^47^. While the exact functions of *yoaB*, *fpaP*(2), *immR*, and *insK1* in LA may vary, their up- and down-regulations highlight their importance to adaptive mechanisms in LA in response to diallyl disulfide.

L-methionine and sodium butyrate had minimal effects on LA gene regulation compared with other treatments. However, it’s important to note that each still had a unique impact. Under L-methionine treatment, the significant up-regulation of *cysM* is noteworthy. This gene plays a role in cysteine biosynthesis, a critical component in the production of proteins and antioxidants^48,49^. The down-regulation of *comGA* and *comGB* is intriguing because these genes are associated with genetic competence in LA, facilitating genetic diversity and adaptability by allowing the uptake and incorporation of external DNA into its genome^44,50^. In contrast, sodium butyrate did not significantly affect gene regulation, with very few log_2_ fold change values above 1.5. Sodium butyrate is a well-studied short-chain fatty acid and a well-known histone deacetylase inhibitor, notably produced in the gut by the microbiota^51,52^. Given that prokaryotes lack histones, the absence of a significant effect on LA might be attributable to this characteristic.

### Modulation of metabolite profiles in *Lactobacillus acidophilus* by dietary bioactive compounds

A comprehensive overview of the primary metabolic profiles for all treated samples compared with wild type is provided in Figure 2. The changes in primary metabolites are indicated by a color key ranging from -0.5 to 0.5.

**Figure 2.**
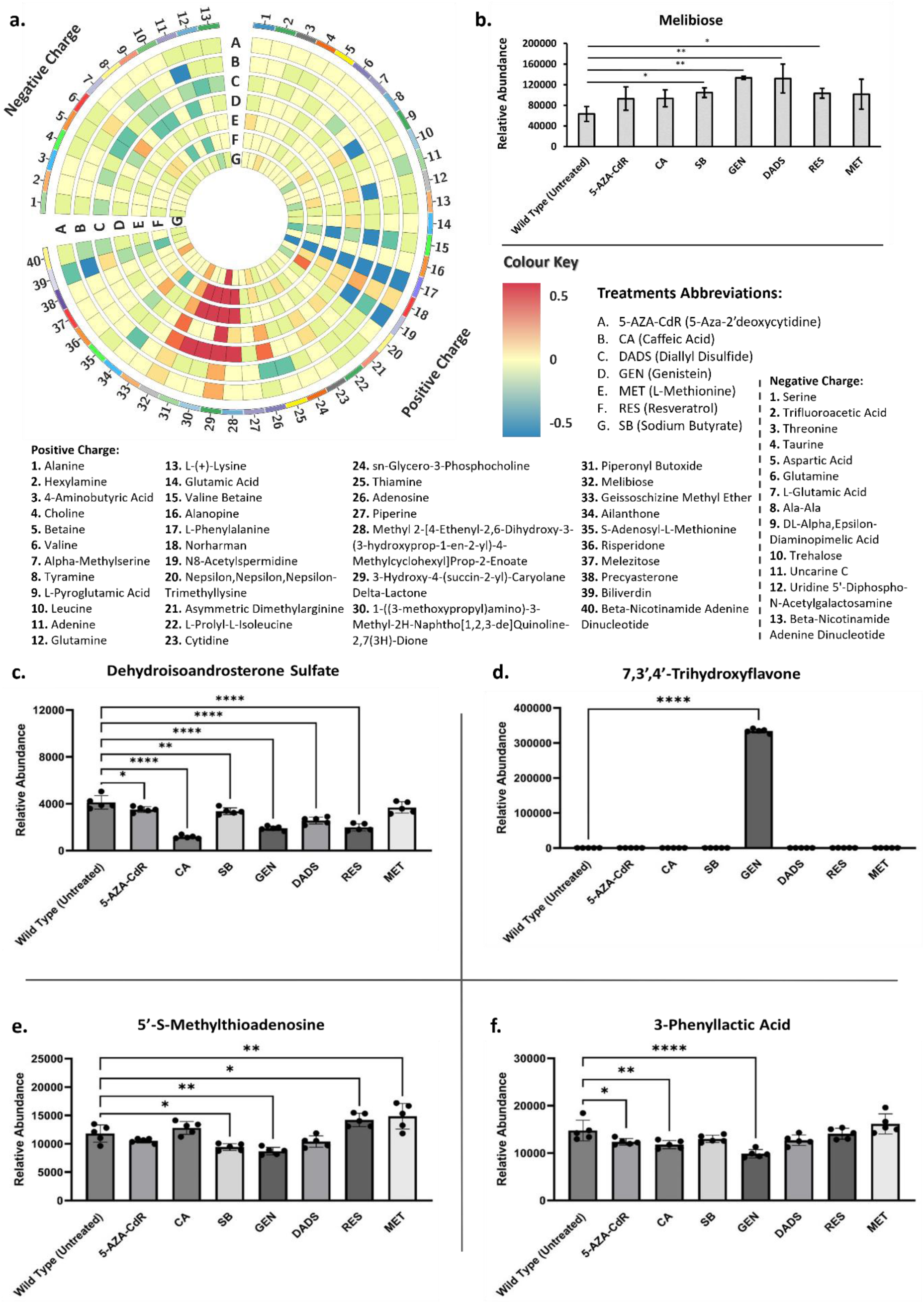
**a.** Circos plot of primary metabolites detected by mass spectrometry (including both positive and negative charge modes) in dietary compound-treated *Lactobacillus acidophilus* ATCC 4356 (LA) compared with the untreated strain. The color key indicates up-regulation (red), down-regulation (blue), and no change (yellow) of each primary metabolite produced by LA differentially treated with the dietary bioactive compounds; **b.** Relative abundance of melibiose in LA when treated with various dietary bioactive compounds, with asterisks marking significant differences (* *p* < 0.05, ** *p* < 0.01). **c-f.** Relative abundance of secondary metabolites when treated with different dietary bioactive compounds, detected by MS1 and MS2 in both negative and positive charge modes. Asterisks denote significant differences (* *p* < 0.05, ** *p* < 0.01, **** *p* < 0.0001). Circular plots represent replicates, with error bars showing standard deviation (S.D.). **Abbreviations:** 5-AZA-CdR: 5-Aza-2’deoxycytidine; CA: Caffeic Acid; DADS: Diallyl Disulfide; GEN: Genistein; MET: L-Methionine; RES: Resveratrol; SB: Sodium Butyrate.

Mass spectrometry identified 53 primary metabolites that had altered concentrations compared with the levels in the untreated strain, with 40 detected in positive and 13 in negative ionization modes, respectively (Figure 2a). Notably, all LA samples treated with dietary bioactive compounds showed significant down-regulation of L-phenylalanine compared with the untreated strain. In addition, each compound induced unique primary metabolic profiles in LA, like that observed during transcriptomic profiling. Unlike the transcriptomic data, however, genistein did not result in the most significant primary metabolic profile differences but did lead to a significant increase in 3-hydroxy-4-(succin-2-yl)-caryolane delta-lactone levels. In contrast, diallyl disulfide, L-methionine, and resveratrol caused more prominent changes in primary metabolic profiles, including significant increases for methyl 2-[4- eth enyl- 2,6- dihydroxy- 3- (3- hydroxyprop- 1- en- 2- yl)- 4- methylcyclohexyl]prop- 2- enoate, 1- ((3- me thoxypropyl)amino)-3-methyl-2h-naphtho[1,2,3-de]quinoline-2,7(3h)-dione, and piperonyl butoxide (Figure 2). Treatment with diallyl disulfide also resulted in the significant reduction in levels of of L- pyroglutamic acid^53^, glutamine^54^, and valine betaine^55^. Each of these compounds play diverse roles in biochemical processes and are essential for various biological functions. The reduced production of these two compounds likely suggests a metabolic shift and reduced nutrient availability in LA in response to diallyl disulfide treatment. Furthermore, compounds such as biliverdin are reported to involved in stress responses. The elevated presence of biliverdin following treatment with diallyl disulfide may indicate that there are notable alterations in primary metabolic pathways as part of the organism’s responses to diverse stresses^56,57^.

Melibiose, a functional carbohydrate and prebiotic^58,59^, was overproduced in LA strains treated with genistein compared with the untreated control (Figure 2b), an observation that aligned with the transcriptomics results. The up-regulation of *melR* and the accumulation of melibiose in genistein-treated LA indicate an adaptive response to genistein exposure. Further investigation is necessary to understand the specific metabolic pathways affected and their implications for LA’s behavior.

Secondary metabolic profiles in LA under different treatments were less complex than their primary metabolic profiles. Four secondary metabolites – dehydroisoandrosterone sulfate (DHEA-S), 7,3’,4’-trihydroxyflavone, 5’-S-methylthioadenosine (5’-S-MTA), and 3-phenyllactic acid – showed significant variation among treatments (Figure 2c-f). Importantly, these metabolites have diverse biological roles in multiple metabolic pathways, many of which are commonly conserved across organisms, and thus have potential health implications for humans.

For example, DHEA-S (Figure 2c) and 7,3’,4’-trihydroxyflavone (Figure 2d) are associated with stress responses. Dehydroisoandrosterone sulfate, the sulphated form of dehydroepiandrosterone (DHEA), plays a role in various physiological processes, including stress response, and has been linked to human mood and overall well-being^60^. Additionally, DHEA-S has been linked with anti-aging effects and increased life expectancy^61^. Yoshida, et al.^62^ reported an inverse correlation between DHEA-S levels and sex-dependent carotid atherosclerosis. All treatments except L-methionine significantly reduced DHEA-S levels compared with the untreated wild type, with changes ranging from *p* < 0.05 to *p* < 0.0001. Notably, compounds which exhibited prominent changes in both transcriptomic and primary metabolomic analyses, such as genistein, diallyl disulfide, and resveratrol, resulted in particularly low DHEA-S levels, potentially impacting stress and aging processes.

On the other hand, 7,3’,4’-trihydroxyflavone has been reportedly associated with reducing oxidative stress, due to its antioxidant properties^63^. Only genistein treatment induced significantly higher levels of this compound (Figure 2d). 7,3’,4’-trihydroxyflavone is also a well-known flavonoid found in various plants^64^ and this observation suggests that under the right conditions, LA may produce or transform bioactive compounds into useful metabolites contributing to its probiotic function.

The third secondary metabolite identified, 5’-S-MTA, participates in the methionine salvage pathway, where it helps in maintaining cellular health^65,66^. Its accumulation increased significantly in strains treated with resveratrol and L-methionine (Figure 2e) indicating that an active methionine salvage pathway that supports cellular health and metabolic competence is present and was activated. Conversely, sodium butyrate and genistein treatments reduced 5’-S-MTA levels, potentially negatively impacting the cellular health and metabolic functions of LA. These observations clearly show that the overall combined impact of metabolite changes in LA for a particular treatment will need to be evaluated. For example, genistein treatment reduced 5’-S-MTA levels but increased melibiose, which is beneficial for human health.

Lastly, 3-Phenyllactic acid levels were found to decrease significantly in LA strains exposed to the positive control chemical 5-aza-2’-deoxycytidine, as well as caffeic acid, and genistein treatments. 3-Phenyllactic acid is a lactic acid variant with a phenyl group attached, which is typically generated during the fermentation process^67^, and is recognized for its antimicrobial properties, inhibiting the growth of various microorganisms including *Salmonella enterica*^67,68^. This observed decrease suggests that the cells may be reallocating resources in response to these compounds, potentially affecting their antimicrobial production^67,68^.

### Epigenetic analysis revealed dietary bioactive compounds influence N4-methylcytosine

The epigenetic status of a bacterial genome is known to have regulatory effects on the transcriptome in bacteria^17,69–71^. Given genistein and resveratrol most influence LA’s transcriptomic profiles, we conducted an in-depth epigenomic analysis focusing on the top 10 up-regulated and top 10 down-regulated genes in genistein-treated LA (Table 1). The study aimed to investigate DNA methyltransferase target motifs in LA and their response to dietary compounds.

**Table 1.** Summary of the top ten significantly up-regulated and down-regulated genes in genistein-treated *Lactobacillus acidophilus* ATCC 4356.

| Gene | Primary accession | Function | Associated domain accession | Regulation* | Protein size (aa) | Modification patterns around promoter (±50bp) |
| --- | --- | --- | --- | --- | --- | --- |
| *melR* | Q5FIG1 | Melibiose operon regulatory protein | PF12833 | 8.860 | 336 | None |
| *LBA1020*<sup>#</sup> | Q5FKA5 | Putative mucus binding protein | PF17965, PF17966, PF04650 | 6.153 | 2311 | Demethylation (4mC) |
| *araC* | Q5FME2 | Arabinose operon control; Helix-turn-helix | PF02311, PF12833 | 5.619 | 274 | Demethylation (4mC) |
| *licT* | Q5FL29 | Transcription antiterminator | PF00874, PF03123 | 3.450 | 281 | None |
| *LBA1031*<sup>#</sup> | Q5FK95 | Protein kinase | - | 2.891 | 198 | Methylation (4mC) |
| *LBA0493*<sup>#</sup> | Q5FLP6 | Aggregation promoting protein | - | 2.759 | 232 | Demethylation (4mC & 6mA) |
| *nagB* | Q5FHT1 | Glucosamine-6-phosphate deaminase | PF01182 | 2.758 | 240 | None |
| *LBA0018<sup>#</sup>* | Q5FMZ9 | GlsB/YeaQ/YmgE family stress response membrane protein | PF04226 | 2.497 | 87 | No promoters predicted |
| *lysA* | Q5FJE6 | Lysin | PF01183, PF03217 | 1.892 | 411 | Methylation (4mC) |
| *LBA0873<sup>#</sup>* | Q5FKP2 | CsbD-like family protein | PF05532 | 1.668 | 58 | Methylation (6mA) |
| *LBA0125<sup>#</sup>* | Q5FMP8 | HTH cro/C1-type domain-containing protein | PS50943 | -3.308 | 270 | Methylation (6mA) |
| *LBA0205<sup>#</sup>* | Q5FMH4 | Heat shock low molecular weight | PF00011 | -2.399 | 142 | Methylation and demethylation (4mC) |
| *groL* | Q93G07 | Chaperonin GroEL | PF00118 | -2.189 | 543 | Demethylation (4mC) |
| *LBA1606<sup>#</sup>* | Q5FIQ3 | DUF4870 | PF09685 | -2.086 | 115 | Demethylation (6mA) |
| *Unnamed* |  | DUF3923 | PF13061 | -2.066 | 75 | Methylation (4mC) |
| *clpE* | Q5FLA7 | ATPase | PF00004, COG0542, PRK10865 | -2.057 | 729 | Methylation (4mC & 6mA) |
| *psiD* | Q5FIQ2 | Phosphatidylserine decarboxylase family protein | PF02666 | -2.028 | 430 | None |
| *proC* | Q5FK14 | Pyrroline-5-carboxylate reductase | PF14748 | -1.979 | 92 | Demethylation (4mC) |
| *scrA* | G1UB46 | Phosphotransferase system | PF00358, PF00367 | -1.949 | 650 | None |
| *proC* | Q5FK13 | Pyrroline reductase | PF14748 | -1.931 | 96 | Demethylation (4mC) |

\*Regulation is represented using log<sub>2</sub> fold change values; positive values indicate up-regulation, and negative values indicate down-regulation; #: UniProt ordered locus name; aa: amino acid units.

Comparative genomic analysis showed that LA wild type (LA-WT), genistein-treated LA (LA-GEN), and resveratrol-treated LA (LA-RES) share identical methylation systems in that 6mA and 4mC modifications were observed throughout the genome. To eliminate the possibility of DNA mutation driving the observed changes, each treated strain was sequenced, and the three genomes were identical, except for three specific point mutations: LA-WT (g.1507125A -> G), LA-RES (g.726428G -> A), and LA-GEN (g.1282181C -> T), none of which are expected to result in phenotypic differences. A total of 17,187, 17,121, and 17,700 genomic positions were methylated (4mC or 6mA) in LA-WT, LA-GEN, and LA-RES, respectively. Analysis of these genomic positions revealed 6mA methylation sites were generally conserved across all the three groups. However, substantial differences in 4mC methylated bases were observed between LA-WT and the treated samples.

Six methylated motifs (one 6mA methylated motif and five 4mC methylated motifs) were identified for each of LA-WT, LA-GEN, and LA-RES and detailed information about these motifs is summarized in Extended Data Table 3. Four motifs (one 6mA and three 4mC) from each strain were selected because of their similarity across the three strains (Figure 3 a (i-iv)). According to Schiffer et al., motifs with a modification quality value (QV) below 50 are likely artifacts^72^. In this study, all identified motifs had a mean modification QV exceeding 130, confirming their authenticity.

**Figure 3.**
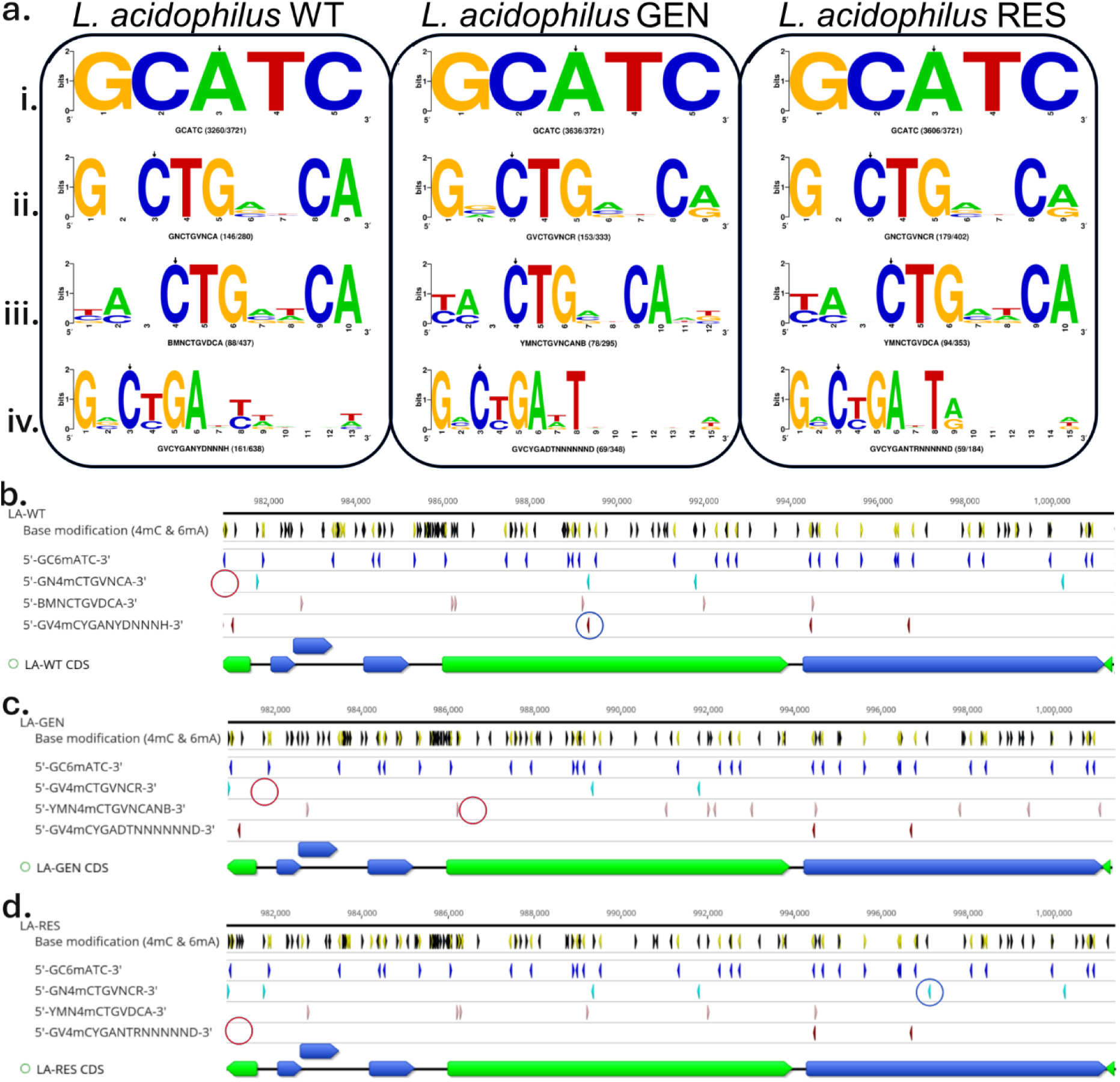
Epigenomic mapping of *Lactobacillus acidophilus* ATCC 4356 (LA) showing **a.** motif logos in untreated LA (*L. acidophilus* WT), genistein-treated LA (*L. acidophilus* GEN), and resveratrol-treated LA (*L. acidophilus* RES) strains. Below each motif logo, the corresponding degenerate motif sequence is shown, along with the total number of its occurrence in the LA genome and the number of times it was modified, expressed as a ratio of modified to total occurrence; **b.** 4mC (black) and 6mA (yellow) base modifications in untreated LA (LA-WT) (track 2) and four motifs in tracks 3 to 6: GC6mATC (blue), GN4mCTGVNCA (cyan), BMN4mCTGVDCA (pink), GV4mCYGANYDNNNH (red); **c.** 4mC (black) and 6mA (yellow) base modifications in genistein-treated LA (LA-GEN) (track 2) and four motifs in tracks 3 to 6: GC6mATC (blue), GV4mCTGVNCR (cyan), YMN4mCTGVNCANB (pink), GV4mCYGADTNNNNNND (red); **d.** 4mC (black) and 6mA (yellow) base modifications in resveratrol-treated LA (LA-RES) (track 2) and four motifs in tracks 3 to 6: GC6mATC (blue), GN4mCTGVNCR (cyan), YMN4mCTGVDCA (pink), GV4mCYGANTRNNNNND (red). These motifs were mapped across the loci containing the coding sequences for the putative mucus binding protein (annotated with navy blue color in the last track of b, c, and d). Motifs modified in one strain but unmodified in others are highlighted with blue circles, while red circles indicate positions where motifs are absent in a strain but modified in others.

One motif, 5′-GC6mATC-3′ (Fig 3 i) containing a 6mA modification, was consistent across all three treatments, with no significant changes between the wild type and treated samples. This Type II restriction-modification (R-M) motif is well-known, with over 1,000 hits on REBASE^73^, and is found in many lactic acid bacteria, including *L. plantarum, L. animalis, L. hordei, Lactococcus lactis, Streptococcus suis,* and *Bifidobacterium longum*^74–76^. Additionally, each of LA-WT, LA-GEN, and LA-RES appeared to have an additional five motifs, all featuring 4mC modifications, which had not been previously reported in the literature. These motifs appear to represent the same DNA sequence recognition sites despite each having slight variations based on the software prediction for each of the strain treatments (Figure 3a). The base modification percentages in motifs with 4mC modifications showed slight variations between the wild type and treated samples (as shown in Extended Data Table 3). However, the modifications in the 5′-GC6mATC-3′ motif remained consistent across all conditions, suggesting that dietary compounds affect 4mC modifications but do not influence 6mA.

For example, a group of motifs (Figure a (ii)) shared most bases, with each treated sample motif differing by one (LA-RES: A -> R) or two (LA-GEN: N -> V and A -> R) bases from the LA-WT (Extended Data Table 3). These motifs were found at varying frequencies in the genomes of the three samples, but the percentage modified in the genome was similar in the treated samples (45.95% in LA-GEN and 44.53% in LA-RES) compared with the wild type (52.14%). Another group of motifs (Figure a (iii)) showed a showed a similar trend (LA-WT: 5′-BMN4mCTGVDCA-3′, percentage modified 20.14%; LA-GEN: 5′-YMN4mCTGVNCANB-3′, percentage modified 26.44%; and LA-RES: 5′-YMN4mCTGVDCA-3′, percentage modified 26.63%).

Epigenomic mapping of loci containing the coding sequences for a putative mucus-binding protein in LA-WT, LA-GEN, and LA-RES is shown in Figure 3b-d. Some motifs, such as the ones in Figure 3 a(i), are conserved across treatments, while others, such as the ones in Figure 3 a(iii), appear to vary depending on the specific dietary compound treatment used. A few 4mC-methylated motifs, such as in Figure 3 a (iv), contain stretches of Ns, indicating a nonspecific spacer region, characteristic of the recognition sequences of Type I and some Type II R-M systems^17^. However, all identified Type I methylases characterized to date, methylate adenine residues, making them 6mA methylases^77^. Therefore, we hypothesize that these motifs might be associated with Type II R-M systems or possibly a new R-M system. Restriction-modification systems are often mobile and variable among bacterial species and strains^17,78,79^. Certain subunits of R-M systems, which recognize specific DNA sequences, contain mobile amino acid sequences that can migrate between different protein domains through recombination of the corresponding DNA sequences. This mobility suggests that R-M systems play a role beyond defense mechanisms; they can establish and sometimes impose epigenetic order on a genome^13^. Methylome diversity acts as a unit for natural selection, with each unique pattern of gene expression corresponding to a distinct set of phenotypes.^17,80^ Except for the 5′-GC6mATC-3′ motif, a search on REBASE did not return any results for the motifs, indicating that these may be novel motifs. This is a noteworthy observation, as it indicates that epigenetics in LA, may be driven most significantly by 4mC modifications rather than 6mA methylation.

The identification of the methyltransferases responsible for these motifs or their corresponding endonucleases was not pursued in this study. Future research will aim to determine whether these methyltransferases are associated with cognate restriction endonucleases or if they are orphan methyltransferases.

The genes that were most highly up-regulated and also those most strongly down-regulated in transcriptomics analysis following genistein treatment of the LA strain, were evaluated for their specific methylation profile (Figure 4 a–f). Differential base modifications were noted around the promoter region and gene body for certain up-regulated genes. For instance, the gene coding for the putative mucus binding protein (Q5FKA5) had a 4mC methylated base around the promoter region in LA-WT but not in LA-GEN (Figure 4b). This pattern was also seen in other up-regulated genes, such as the gene coding for the arabinose operon control (*araC)* (Figure 4c). Conversely, a reverse trend was observed in up-regulated genes such as those coding for protein kinase (Q5FK95) and lysin (Q5FJE6) (Table 1). While methylation changes in a promoter region might be expected to have a significant effect on gene expression, we found differences in the gene body were more pronounced, with more instances of methylation and demethylation in response to treatment identified, which may also contribute to gene expression. Interestingly, there was also a mix of methylation and demethylation in the promoter region and gene body of downregulated genes (Figure 4d-f & Table 1). For instance, both a methylation and demethylation change were observed in the promoter region of the gene coding for the heat shock protein (Q5FMH4, Figure 4e) and the chaperonin protein (Q93G07, Figure 4f), respectively.

**Figure 4.**
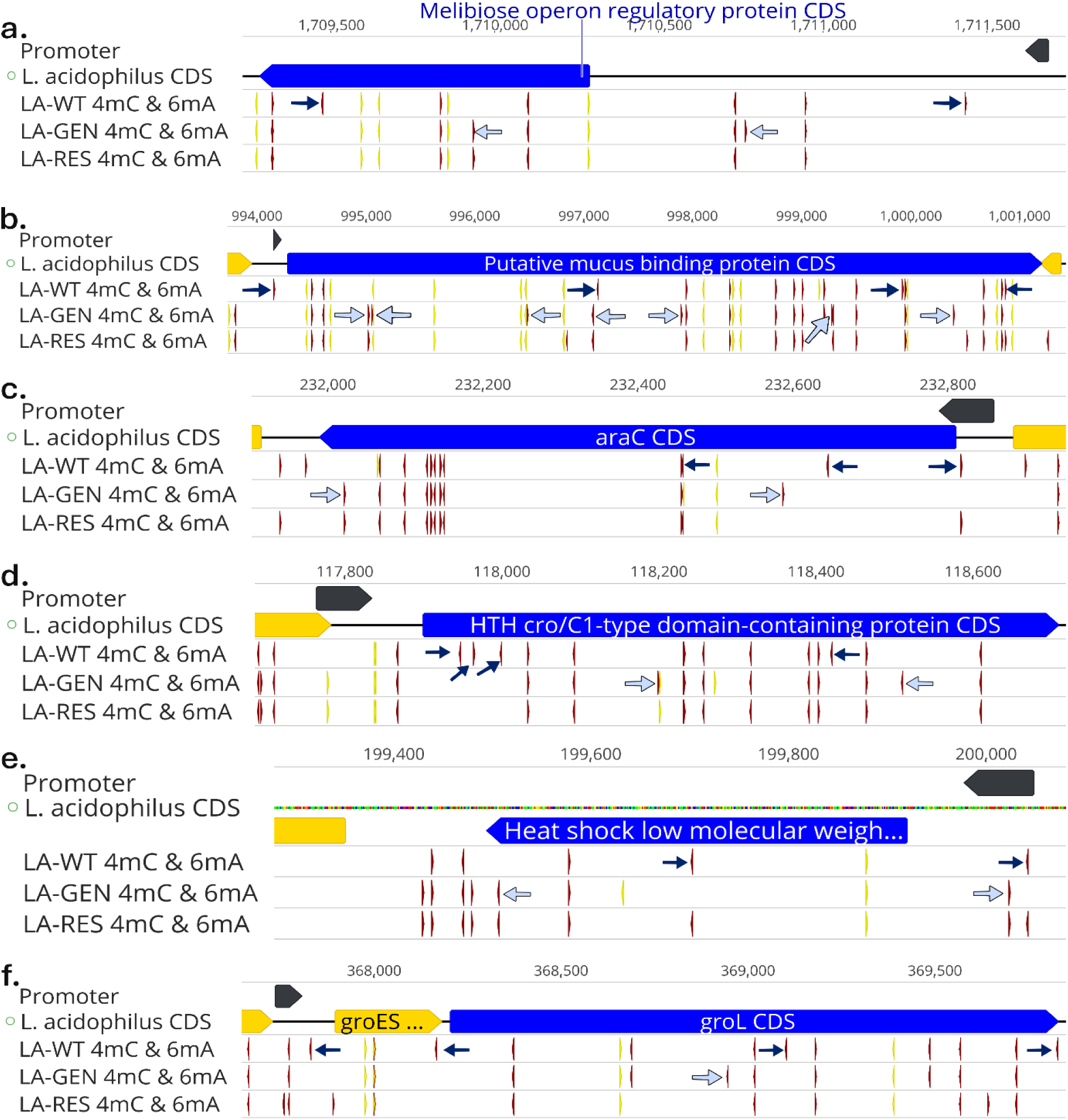
Map of the top three significantly up-regulated (**a-c**, highlighted in blue) and down-regulated (d-f, highlighted in blue) genes in genistein-treated *Lactobacillus acidophilus* ATCC 4356 (LA) (LA-GEN) showing the effect of the treatment on the methylation patterns around the promoter region and within the gene body. The first track in each panel (**a-f**) shows the predicted promoter region (black arrow) for the gene. The second track contains the coding sequences in LA genome (blue and yellow arrows). The third, fourth and fifth tracks contain 4mC (in red) and 6mA (in yellow) modification regions in untreated LA (LA-WT), LA-GEN, and resveratrol-treated LA (LA-RES), respectively. The 4mC-modified base(s) in LA-WT but unmodified in LA-GEN are indicated by a dark blue arrow in track three of panels **a-f**. Conversely, 4mC-modified base(s) in LA-GEN but unmodified in LA-WT are annotated with a light blue arrow in track four of panels **a-f**.

To confirm our findings, we conducted epigenomic analyses on different species. We selected two up-regulated and two down-regulated genes in LA with genistein treatment that have homology with *Lacticaseibacillus paracasei* ET22 and compared LA with genistein and resveratrol treatments to *Lacticaseibacillus paracasei* ET22 with the same treatments (Extended Data Fig 2). We observed conservation of 6mA sites regardless of treatment, but changes were observed for 4mC depending on the treatment in a manner similar to that for LA; however, the epigenetic profiles of the two organisms differ, so the epigenetic impact on gene expression is likely to be strain dependent^18^. Additionally, we analyzed another strain, *Lacticaseibacillus paracasei* K56, treating it with caffeic acid and resveratrol. Two up-regulated and two down-regulated genes in LA that have homology with *Lacticaseibacillus paracasei* K56 were examined. The dietary compounds had a similar effect on 4mC but no significant effect on 6mA, as observed in LA (Extended Data Fig 3).

Since these compounds do not appear to uniformly affect all genes, caution should be exercised when interpreting the effect of methylation or demethylation on individual gene expression, therefore, warranting further investigation.

### Enhancing probiotic functionality in *Lactobacillus acidophilus* through dietary bioactive compounds

The findings from transcriptomic, metabolomic and epigenetics data provide insights into the potential of dietary compounds for microbial metabolic engineering. They highlight the importance of considering the unique responses of individual microbial strains to improve the probiotic functionality of LA.

Mucins or mucin-like proteins, expressed by epithelial cells in the vagina, digestive tract, and respiratory system, play a critical role in mucosal immunity. Bacterial adhesion through binding domains in the mucosal layer facilitates colonization. Mucin-binding proteins are essential for bacterial adhesion to host mucus, contributing to the bacteria’s ability to adhere and colonize surfaces, and are well-characterized among *Lactobacillus* species. Recently, Wei et al. (2024) observed that genes encoding bacterial adhesion components, including mucin-binding genes (*muc_B2/mucBP/mucBP_2*) and cell wall anchor genes of Gram-positive bacteria, were under positive selection. These genes are crucial for the host-microbe interface at epithelial surfaces throughout the human body. Wei et al. demonstrated that multiple mucin-binding genes in several *Lactobacillus* species experienced repetitive positive selection pressure in the vaginal microbial environment^81^.

The volcano plots illustrate *p*-values and log_2_ fold changes between the following treatment pairs: untreated LA (LA-WT) and genistein-treated LA (LA-GEN), LA-WT and resveratrol-treated LA (LA-RES), and LA-RES and LA-GEN. Red dots in panels d and f represent up-regulated proteins (log_2_ fold change ≥ 1.0; *p*-value < 0.05) in LA-GEN compared with LA-WT and LA-RES, respectively. Yellow dots in panel e indicate up-regulated proteins in LA-RES compared with LA-WT. Blue dots in panels d and e, and yellow dots in panel f, indicate down-regulated proteins (log_2_ fold change ≤ -1.0; *p*-value < 0.05) in the same comparisons. Proteins not meeting these criteria are shown as grey dots. The vertical purple lines represent the *x* intercept (log_2_ fold change -1.0, 1.0), while the horizontal grey line represents the *y* intercept (-log_10_(0.05)). Annotated proteins correspond to genes among the top up-regulated and down-regulated genes in the transcriptomic analysis, and also detected in the proteomics data.

A comparative analysis of the responses of LA to genistein and resveratrol treatment is shown in Figure 5. The treatments distinctly separated from the wild type in the PCA across all three analyses (transcriptomic, metabolomic, and proteomic) as shown in Figure 5 (a-c). The pattern of separation between the treatment groups varies between the different analyses. Our study found that genistein treatment up-regulated a putative mucus-binding gene, increasing the associated protein production (Figure 5 g-j). Interestingly, proteomic analysis demonstrated three other mucus-binding proteins were also produced in higher quantities (Figure 5), although no significant expression changes were observed in transcriptomic data, potentially indicating that the epigenomic changes are originating elsewhere in the genome and impacting for instance on global regulators^82^. This suggests genistein may enhance mucin-binding, potentially through epigenetic regulation as observed in Figure 4.

**Figure 5.**
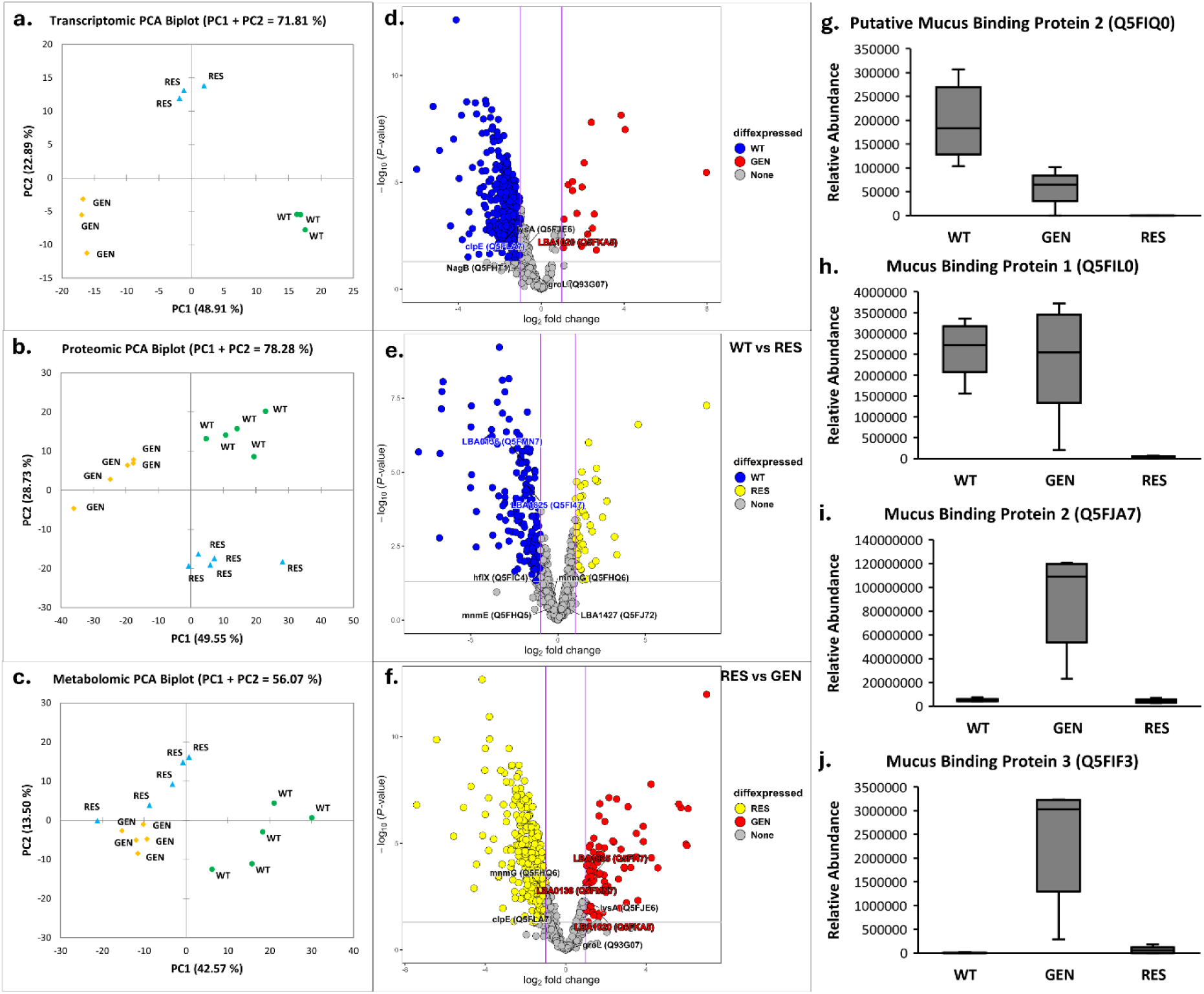
**a-c**. Principal Component Analysis (PCA) score plots generated from transcriptomic, proteomic and metabolomic results of three different *Lactobacillus acidophilus* ATCC 4356 (LA) groups treated with dietary bioactive compounds: wild type (WT), genistein (GEN), and resveratrol (RES); **d-f.** Volcano plots of differential protein analysis in the three groups. **d.** WT vs GEN annotations: *nagB* (Q5FHT1), l*ysA* (Q5FJE6), *groL* (Q93G07), *clpE* (Q5FLA7) and *LBA1020* (Q5FKA5); **e.** WT vs RES annotations: *hflX* (Q5FIC4), *mnmG* (Q5FHQ6), *mnmE* (Q5FHQ5), *LBA0136* (Q5FMN7), *LBA1825* (Q5FI47) and *LBA1427* (Q5FJ72); **f.** RES vs GEN annotations: *mnmG* (Q5FHQ6), *lysA* (Q5FJE6), *clpE* (Q5FLA7), *groL* (Q93G07), *LBA1825* (Q5FI47), *LBA0136* (Q5FMN7) and *LBA1020* (Q5FKA5). **g-j.** Relative abundance of one putative mucus binding protein and three mucus binding proteins produced by LA when treated with dietary bioactive compounds, identified by proteomic analysis with Tukey pairwise mean comparisons (*P* < 0.05). **g.** Putative Mucus Binding Protein 2 (Q5FIQ0): WT (a), GEN (b), RES (b); **h**. Mucus Binding Protein 1 (Q5FIL0): WT (a), GEN (a), RES (b); **i.** Mucus Binding Protein 2 (Q5FJA7): WT (b), GEN (a), RES (b); **j.** Mucus Binding Protein 3 (Q5FIF3): WT (b), GEN (a), RES (b).

A second example from genistein treatment includes significant up-regulation of melibiose production. Melibiose (α-d-galactopyranosyl-(1→6)-α-d-glucopyranoside) is a reducing disaccharide composed of galactose and glucose linked via an α-1,6 bond. It has gained increasing attention for its beneficial properties with the benefit that appropriate intake of melibiose fosters the growth of bifidobacteria, improving stool consistency. Dietary melibiose effectively suppresses the Th2 immune response and enhances oral tolerance^83^. Its potent prebiotic properties make it a valuable additive in functional foods and pharmaceuticals. However, melibiose is challenging to obtain through conventional methods and is primarily produced via enzymatic trans-glycosylation reactions using raffinose and lactose or galactose as substrates^83,84^. This study is the first to demonstrate significant melibiose production through treatment with genistein (Figure 2b) without genetic modification or enzymatic reactions, highlighting the potential for using dietary bioactive compounds to enhance production of beneficial metabolites. A third example from genistein treatment includes an increase in 7,3’,4’-trihydroxyflavone (Figure 2). Although metabolomics studies have identified many microbial metabolites of polyphenols in systemic circulation, these compounds have yet to be thoroughly investigated^85^. This relatively rare flavone, first isolated from *Trifolium repens* and *Medicago sativa* in the 1960s, is known for its bioactive properties, including anticancer, antioxidant, and anti-inflammatory activities^86–88^. It may just be a byproduct of LA metabolism of genistein but provides evidence that natural compounds can be synthesized using this method.

In the future, by targeting specific genes and metabolic pathways through epigenetic modulation, it is possible to optimize the health benefits of these probiotics. This study provides evidence for beneficial metabolites and natural product synthesis using widely consumed probiotic like LA and dietary bioactive compounds like genistein.

## Discussion

This study provides a comprehensive analysis of the transcriptional, metabolic, proteomic, and epigenetic effects of specific dietary bioactive compounds on LA. Each dietary compound uniquely influenced LA, indicating a high level of specificity in how these compounds modulate gene expression and metabolism. Notably, the study identifies the potential role of 4mC in LA gene regulation, an area relatively unexplored in bacterial epigenetics.

The differential expression patterns observed following LA treatment, particularly with genistein, demonstrate significant shifts in gene regulation (Figure 1). The up-regulation of genes related to carbohydrate metabolism, such as those involved in melibiose and maltose utilization, indicates that genistein can enhance the metabolic capabilities of LA towards certain sugars. Additionally, the observed down-regulation of genes associated with galactose and lactose metabolism indicate a possible trade-off or reprogramming of metabolic pathways in response to genistein.

Metabolomic analysis supported the transcriptomic findings, showing distinct metabolic profiles for LA treated with different compounds (Figure 5). Modulation of metabolic profiles in LA by these compounds, particularly the significant up-regulation of metabolites like melibiose and 7,3’,4’-trihydroxyflavone, show the potential for specific dietary compounds exposure to manipulate microbial metabolism in beneficial ways (Figure 2). Furthermore, the specific metabolomic signatures induced in LA by each compound, provides evidence that tailoring of gut microbiota functions may be feasible through directed dietary modulation and underscores the need to consider individual microbial responses when designing dietary interventions or developing probiotic-based therapies.

Our observation of the minimal impact of sodium butyrate on gene regulation in LA, despite its known activity for influencing eukaryotic epigenomes, suggests that the mechanisms by which it influences bacterial gene expression may differ significantly from those in eukaryotic systems, possibly due to the absence of histones in prokaryotes.

Methylation of DNA base pairs in promoter regions significantly influences gene expression, often reducing or completely blocking transcription and silencing the gene^89^. Genes with gene-body methylation typically show moderate expression across many tissue types^89^. These methylation-mediated alterations in gene expression can enhance an organism’s fitness. For example, DNA methylation facilitates the colonization of host environments by bacteria such as *Helicobacter pylori*^90^ and enables *Lacticaseibacillus paracasei* to regulate carbohydrate metabolism, improving growth under nutrient-deficient conditions^91^.

Methylation of the promoter region of a key gene, like the one for a mucus-binding protein that helps lactobacilli stick to mucosa, occurs in untreated LA but not in genistein-treated LA. This suggests that genistein treatment might boost the probiotic potential of LA by activating this gene. Zhao et al. (2023) proposed that 6mA modifications are involved in carbohydrate metabolism in *Lacti. paracasei*^91^. By contrast, this present study indicates that 4mC modifications may also play a role in carbohydrate metabolism and are influenced by dietary compounds, whereas these compounds might not affect 6mA modifications. This alteration in carbohydrate metabolism was clearly observed in the genistein treatment on LA (Table 1, Figure 1, and Extended Data Fig 1).

This new approach termed “Nutrifermentics”^8^ offers a novel, non-GMO approach to microbial metabolic engineering by leveraging the natural regulatory effects of dietary compounds. This method has potential to significantly impact the development of functional foods and probiotic supplements aimed at enhancing human health. Once this research direction is validated by other chemical biologists, its simplicity and non-permanence could facilitate its adoption across various research settings and industries. The technique does not require sophisticated equipment, and the temporary nature offers advantages to commercial strain manufacturers, allowing them to retain intellectual property or increase regular sales.

In conclusion, this study illustrates the diverse and gene-specific effects of dietary bioactive compounds on LA, providing insights into their potential for microbial metabolic engineering. The findings highlight the importance of considering the unique responses of individual microbial strains and the potential for dietary bioactive compounds to enhance probiotic functionality. Future research should focus on elucidating the role of 4mC in gene regulation and its possible involvement in novel restriction-modification systems in bacteria. The effects of dietary compounds on 4mC as observed in this present study, should be further explored in eukaryotic and human cell lines to determine their influence on 4mC in these systems. Finally, the compounds evaluated in this study, along with other dietary bioactive compounds, need to be tested at various concentrations and combinations, for their impact on different gut microbiota strains.

## Methods

### Growth conditions of Lactobacillus acidophilus

*Lactobacillus acidophilus* ATCC 4356 (LA) was used as the model strain for all experimental analyses. This strain was sourced from the Institute of Environmental Science and Research (ESR), New Zealand. Cultivation was done anaerobically in De Man, Rogosa and Sharpe (MRS) Broth (Fort Richard Laboratories Ltd, New Zealand) in 15 mL Falcon tubes, with some cultures supplemented with dietary bioactive compounds. Cultures were incubated at 37 °C for 15 h, with continuous shaking at 120 rpm. To systematically investigate the effect of dietary bioactive compounds on LA, we selected previously described compounds^8^, including caffeic acid, diallyl disulfide, genistein, L-methionine, resveratrol, and sodium butyrate. Table 2 below lists the seven dietary bioactive compounds and their concentrations used, with 5-aza-2’-deoxycytidine as the positive control. Each compound’s concentration was optimized to minimize its impact on LA growth while inducing significant gene expression shifts due to epigenetic changes. These compounds were obtained from MedChemExpress (MCE), New Jersey, USA.

**Table 2.** Dietary bioactive compounds used in this study and their concentrations.

| Compound | Abbreviation | Concentration |
| --- | --- | --- |
| 5-Aza-2'-deoxycytidine | 5-AZA-CdR | 1 mM |
| Caffeic Acid | CA | 2.5 mM |
| Diallyl Disulfide | DADS | 1.5 mM |
| Genistein | GEN | 500 µM |
| L-Methionine | MET | 1 Mm |
| Resveratrol | RES | 500 µM |
| Sodium Butyrate | SB | 5 mM |

### DNA methylation analysis

#### DNA extraction

The bacterial DNA samples from *Lactobacillus acidophilus* ATCC 4356 (LA), *Lacticaseibacillus paracasei* K56 and *Lacticaseibacillus paracasei* ET22 were extracted using the GenElute™ Bacterial Genomic DNA Kit (Sigma-Aldrich, Missouri, USA), following the manufacturer’s instructions. Their quality was assessed using the DeNovix DS-11 instrument (DeNovix Inc., Delaware, USA), and their concentrations were determined by the Qubit™ dsDNA Quantification (Broad Range) Assay Kit (Invitrogen™, Massachusetts, USA).

#### DNA sequencing and analysis

DNA samples were sent to Azenta Life Sciences (Suzhou, China) for Pacific Biosciences (PacBio) Single Molecule, Real-Time (SMRT) sequencing on the PacBio Sequel system. Azenta performed quality control, data correction and assembly using Canu (v1.7)^92^ and were corrected with Pilon (v1.22)^93^ using prior Illumina sequence data.

The assembled bacterial genomes were annotated with the Bakta annotation pipeline (v1.9.1)^94^ and the Rapid Annotation using Subsystem Technology (RAST) platform^95^. Gene domains were annotated with the DomainAnnotation (v1.0.10) tool on KBase^96^, and the associated domain accession of a gene retrieved from its entry on UniProtKB^97^. PacBio SMRTlink software detected base modifications and calculated motif analysis statistics for the genomes. Degenerate motifs obtained from PacBio SMRT sequencing analysis were then searched on Geneious Prime® 2024.0.5 (https://www.geneious.com) using the *Find Motifs* function based on the EMBOSS (v6.5.7) tool, *fuzznuc*^98^. The bases were searched on both forward and reverse strands, with no mismatches allowed. The consensus sequence of each of the degenerate motifs was then used to make a sequence logo using the WebLogo server (v3.7.12)^99^. The location of the bases identified from the degenerate motifs was confirmed to have undergone a base modification event by annotating with the raw motif signals obtained by SMRT sequence analysis.

Ten significantly up-regulated and down-regulated genes between genistein-treated LA and untreated control, as well as resveratrol-treated LA, were selected for further in-depth analysis. These genes had motifs with base modifications with an identification Qv of at least 20 (1% error), and an up-regulation of log_2_ fold > 1.5 or down-regulation of log_2_ fold < -1.5 with genistein and an up-regulation of log_2_ fold > 1.2 or down-regulation of log_2_ fold < -1.2 with resveratrol. Promoter regions were predicted using the ProPr: Prokaryote Promoter Prediction webserver (v2.0) (accessed on 11-04-2024), which is part of the PEPPER toolbox^100^. Weak promoters (with *min10score* < 5) were excluded. When multiple promoter sequences were predicted for the same gene, promoter sequences with the highest *score*, *min10score*, and correct orientation were selected. Differential modification around the promoter was determined to be within a conservative threshold of 50 bp from the promoter sequence. Demethylation was defined as a base modification present in LA wild type but absent in the treated samples, while methylation was defined as the presence of a base modification in the treated samples but not in the wild type.

To investigate genistein’s effects on similar genes in *Lacticaseibacillus paracasei* ET22, top ten significantly up-regulated and down-regulated gene sequences in LA-GEN were searched against a custom protein database of genistein-treated *Lacticaseibacillus paracasei* ET22 (ET-GEN) using BLASTX (v 2.16.0+) on Geneious Prime®. The search parameters included a maximum e-value of 1e-10, a word size of 3, low complexity filter on, and a maximum of 10 hits. Results were sorted by grade, a percentage calculated by Geneious Prime® by combining the query coverage, e-value, and identity values for each hit with weights of 0.5, 0.25, and 0.25, respectively. Modification patterns around gene bodies and promoter regions of three up-regulated and down-regulated genes with the highest grade in ET-GEN were then investigated.

### RNA transcriptomics analysis methods

#### Assessment of RNA integration using Fragment Analyzer

RNA samples from *Lactobacillus acidophilus* ATCC 4356 (LA) were purified using the RiboPure™ RNA Purification Kit for yeast (Invitrogen™, Massachusetts, USA), following the manufacturer’s instructions, and the RNA integration was assessed using the 5300 Fragment Analyzer System (Agilent Technologies, California, USA) with the Agilent DNF-471 RNA Kit (15 nt), following the kit protocol. The RNA exhibited high integrity for sequencing, with A260/280 >2.0, RQN >6.5 and 23s/16s >1.0.

#### RNA stabilization and sequencing

RNA samples (>3 µg) were transferred into RNA Stabilization Tubes from Azenta Life Sciences (Massachusetts, USA), air-dried in a biosafety hood for 24 h, and shipped to Azenta’s NGS Laboratory (Suzhou, China) at room temperature. Library preparation and transcriptomic sequencing were conducted by Azenta Life Sciences on the Illumina Novaseq platform (Illumina, California, USA), in a 2x150bp paired-end (PE) configuration with triplicate RNA samples for each treatment.

#### RNA bioinformatic analysis

Azenta Life Sciences conducted the bioinformatic analysis, using the reference genome from LA’s *de novo* DNA sequencing results performed on the PacBio platform.

##### Quality control

To remove technical sequences such as adapters, PCR primers, or low-quality bases (Phred score < 20), pass-filter data in fastq format underwent processing using Cutadapt (version 1.9.1) with the following parameters: phred cutoff: 20, error rate: 0.1, adapter overlap: 1 bp, minimum length: 75, proportion of N: 0.1.

##### Mapping

The reference genome sequence was indexed and clean data aligned to the reference genome using Bowtie2 (v2.2.6).

##### Expression analysis

Transcripts in fasta format were generated from known gff annotation files and indexed. Gene expression levels were estimated from the pair-end clean data using HTSeq (v0.6.1p1) with the indexed file as the reference.

##### Differential Expression Analysis

Differential expression was analyzed using the DESeq2 Bioconductor package, which is based on the negative binomial distribution. After adjustment by the Benjamini and Hochberg method to control the false discovery rate, genes with a Padj value < 0.05 were considered differentially expressed^101^.

##### GO and KEGG enrichment analysis

GOSeq (v1.34.1) was employed to identify Gene Ontology (GO) terms annotating a list of enriched genes with a significant padj value < 0.05. Additionally, topGO was used to visualize the Directed Acyclic Graph (DAG). For KEGG enrichment, in-house scripts were used to identify significant differentially expressed genes in KEGG pathways.

### Proteomics methods

#### Bacterial sample preparation

*Lactobacillus acidophilus* ATCC 4356 cultures were divided into three groups: two treatment groups (LA-GEN with 500 µM genistein, LA-RES with 500 µM resveratrol) and an untreated control (LA-WT), each with five biological replicates. Overnight LA cultures were inoculated at a 1:100 ratio into 15 mL of Lactobacilli De Man-Rogosa-Sharpe (MRS) broth (Difco, USA). After 15 h incubation at 37 °C with shaking at 120 rpm, cultures were adjusted to an OD_600_ of 1.0 with sterile MRS broth to standardize across all samples. Bacterial cells were harvested by centrifugation at 8000 × g for 5 min at 4 °C, with pellets were washed three times in phosphate-buffered saline (PBS) (Invitrogen, USA, pH 7.4).

#### Protein extraction

The bacterial pellet was resuspended in 200 µL of lysis buffer (100 mM Tris, 50 mM dithiothreitol, and 2% sodium deoxycholate (SDC)). After vortexing, 100 µL of lysate was transferred into a new microcentrifuge tube. The samples were sonicated thrice for 4 s each on ice. Samples were centrifuged at 14,000 × g for 30 min at 4 °C. Protein precipitation was performed with CHCl_3_/MeOH precipitation, and the pellet was resuspended in 100 µL of 0.1 M ammonium bicarbonate (AMBIC). Protein concentration was determined using an Implen spectrophotometer.

Subsequently, 100 µg of protein was transferred into a protein LoBind® tube (Eppendorf) and alkylated by adding 2 µL of 200 mM iodoacetamide (IAM) and incubation at 25 °C for 45 min. Proteins were then digested with Trypsin/Lys-C (1:50 enzyme:protein) at 37 °C with shaking overnight. Digestion was halted the next day and SDC was precipitated by adding 4 µL of formic acid (FA). Samples were centrifuged at 14,000 x g for 30 min at 4 °C, and the clear supernatant was transferred into a new protein LoBind® tube. The supernatant was cleaned with C18 tips according to the manufacturer’s instructions (Thermo Scientific, Waltham, MA, USA) and dried in a vacuum centrifuge. Dried peptides were resuspended in 50 µL of 0.1% FA, quantified with an Implen spectrophotometer, and 250 ng was injected for Liquid chromatography-mass spectrometry (LC-MS) analysis.

#### LC-MS parameters

Samples were analyzed using a NanoElute 2 LC system (Bruker Daltonics) coupled to an Impact II Q-TOF mass spectrometer equipped with a CaptiveSpray source (Bruker Daltonik, Bremen, Germany). For each sample, 250 ng of the sample was loaded on a Bruker PepSep Fifteen (150 mm x 75 µm, 1.9 µm) analytical column. The reverse phase elution gradient was: 2% to 24% in 70 min, 24% to 50%B in 15 min, 50% to 95% in 2 min, maintaining at 95%B for 3 min, total 90 min at a flow rate of 400 nl/min. Solvent A was LCMS-grade water with 0.1% FA; solvent B was LCMS-grade acetonitrile with 0.1% FA. The LC was interfaced with a captive spray ion source (3.0 L/min dry gas, 1500 V) to an Impact II quadrupole-time-of-flight (Q-TOF) (Bruker Daltonics) mass spectrometer. To profile protein expression patterns, the analytes were detected via MS-only mode in positive ion mode, with a mass range between 150–1800 m/z and a sampling rate of 2 Hz. To link the expression levels with identifications, pooled samples per treatment were run via LC-MS/MS with data-dependent auto-MS/MS mode with the following settings: the same LC parameters as described before, a full scan spectrum, with a mass range of 150-1800 m/z, was followed by a maximum of ten collision-induced dissociation (CID) tandem mass spectra (350 to 1800 m/z) at a sampling rate of 2 Hz for MS scans and 2 to 32 Hz for MS/MS. Precursors with charges 2+ to 5+ were preferred for further fragmentation and a dynamic exclusion of 20 sec was set. Following the LC-MS run, the QTOF data were further analyzed with Compass DataAnalysis 6 software (Bruker Daltonics) to evaluate the LC chromatogram and the overall quality of both MS1 and MS2 spectra.

#### Protein identification and quantification

Raw LC-MS data was analyzed with PEAKS X Pro software (Bioinformatics Solutions Inc., Waterloo, ON, Canada). The raw data was refined by a built-in algorithm. The proteins/peptides were identified with the following parameters: precursor mass error tolerance of 10 ppm, fragment mass error tolerance of 0.05 Da, and the *Lactobacillus acidophilus* reference proteome database (v2024.05, 1859 sequences). Trypsin/LysC was specified as digestive enzyme in semi-specific digest mode, and up to three missed cleavages were allowed. Carbamidomethylation of cysteine was set as a fixed modification and oxidation (M), and deamidation (NQ) as a variable modification. Optimized Peaks PTM settings included additional variable modifications: Acetylation (N-term), Pyro-glu from Q, Pyro-glu from E, Acetylation (N-term), dehydration, Carbamylation, and Carbamidomethylation (DHKE, N-term), Phosphorylation (ST), Amidation (N-term), Formylation (K,M-term), Methylation (DEHST). A maximum of 3 post-translational modifications (PTMs) per protein were permitted. False discovery rate (FDR) estimation was made based on decoy-fusion. An FDR of < 5% with a peptide hit threshold was used for confident protein identification. At least one unique peptide per protein was required for both identification and quantification purposes.

#### Label-free quantification

To relatively quantify the protein expression levels, label-free quantification (LFQ) was performed using the quantitation node of Peaks Studio X Pro software. Relative abundance levels between the proteins in all samples were compared. The following parameters were included: a mass tolerance error of 15 ppm and a retention time (RT) shift tolerance of 2 min was allowed. To determine the relative protein and peptide abundance in the RT aligned samples, peptide feature-based quantification was performed. Relative comparison between samples was based on the area under the curve and to get this cumulative area for each protein, only unique peptides that are assigned to a particular protein were selected. Data were exported for further statistical analysis.

#### Data visualization

Partial least square-discriminant analysis (PLS-DA) plots were generated using the mixOmics package^102^. Proteins were compared between groups by fitting a linear model to the individual protein abundances, using empirical Bayes to accurately estimate the variance (using limma^103^). Proteins were deemed differentially expressed if log2fold change was >1, and the multiple test adjusted *p*-values < 0.05. All graphs and analyses were made using custom scripts developed in R (version 4.3.1).

### Metabolomics methods

#### Lysate extraction

Five mL of freshly grown LA (OD_600_ of 1.0 ± 0.2) were quenched with pre-cooled MeOH/ddH2O (60:40) containing 10 mM ammonium acetate at a 1:4 ratio. The mixture was placed at -80°C for 2 minutes, then centrifuged at 5,000 ’g for 5 min at -10 °C. Intracellular metabolites were extracted by re-suspending the cell pellet in 1 mL of 80% EtOH. The suspension was heated to 80 °C for 2 min with 10 s of vigorous vortexing in between, followed by another 2 minutes of heating. Finally, the mixture was centrifuged at 16,000 ’g for 5 min at -10 °C (Thermo Scientific™ Sorvall™ Legend™ XTR Centrifuge, Thermo Fisher Scientific, Massachusetts, USA). The supernatant, containing intracellular metabolites, was aliquoted and stored at -80 °C until Liquid Chromatography with tandem mass spectrometry (LC-MS/MS) analysis was carried out by the Proteins and Metabolites Team at AgResearch (Lincoln, New Zealand). Each treatment included five biological replicates.

#### Sample preparation

Lysates were stored at -80 °C and thawed overnight at 4 °C. Each 200 µL sample was mixed with 800 µL of ice-cold MeOH/ddH2O (1:1), shaken in a TissueLyser (Qiagen, Venlo, Netherlands) and centrifuged at 4°C for 20 min, at 14000 ’g. Two 200 µL aliquots of the supernatant were extracted: one for sample analysis and one for pooled QC. These samples, along with 200 µL of the pooled QC samples, dried in a vacuum concentrator (Vacuumbrand, Wertheim, Germany) and reconstituted in MeCN:ddH2O (1:1), with L-Tyrosine-d2 as an internal standard to monitor sample degradation.

#### Chromatography

Samples were analyzed using a Nexera X2 ultra high-performance liquid chromatography (UHPLC) system (Shimadzu, Japan), which comprised a SIL-30AC autosampler linked to an LCMS-9030 quadrupole time-of-flight (Q-TOF) mass spectrometer (Shimadzu, Japan) equipped with an electrospray ionization source. A 2 µL sample was loaded onto a normal-phase Ascentis® Express HILIC UHPLC column (2.1 x 100 mm, 2 µm particle size; Sigma-Aldrich, Missouri, USA) and eluted over a 20-minute gradient at 30°C at a flow rate of 400 µL/min. The mobile phase solvent A was 10 mM ammonium formate in water, while solvent B was acetonitrile with 0.1% FA. The solvent gradient program initiated with 97% solvent B from 0 to 0.5 min, decreased to 70% within 11.5 minutes, further decreasing to 10% from 11.5 to 13.5 min, held at 10% for 1.5 min, increased to 97% B within 1 min, and maintained that concentration until the end of the elution process.

#### Mass spectrometry

Full scans (m/z 55-1100) and MS/MS scans covering windows from m/z 20 were performed in both positive and negative ionization modes. The set up included 42 events with a loop time of 0.85 s. Spray voltage was maintained at 4.0 kV for positive ionization mode and -3.0 kV for negative ionization mode. Collision energy (CE) was set at 23 ± 15 V. The ion source operated under optimal conditions: nebulizing gas flow at 3.0 L/min; heating gas flow at 10.0 L/min; interface temperature at 300 °C; drying gas flow at 10.0 L/min; desolvation line temperature at 250°C, and heat block temperature at 400 °C.

#### Batch sequence

Samples were analyzed in a single batch, beginning with positive ionization mode. The sequence started with five blanks, followed by an external standard (Amino acid standard, A9906, Sigma, USA) to verify system performance. QC (Quality Control) pooled samples were run, followed by the actual samples in a randomized order with QC samples interspersed once for every 8 samples. The same sequence was followed for the negative ionization mode.

#### Data analysis

Raw data files (.lcd) were converted to the mzML file format using LabSolutions software (Version 5.99 SP2, Shimadzu, Japan). MS-Dial was employed for peak detection, MS2 deconvolution, alignment of samples, and compound identification^104^. Appropriate adducts were chosen for both positive and negative ionization modes.

Blank data was subtracted from all samples, ensuring that the sample’s maximum ratio to the blank average did not exceed 5. To correct for signal variation, a locally weighted scatterplot smoothing (LOWESS) method was applied during data pre-processing. Only features with metabolite IDs that had MS/MS matches or matched the m/z values (without MS/MS) were exported, along with their respective normalized peak areas, for subsequent statistical analysis. Statistical analysis of the normalized data was carried out using MetaboAnalyst 5.0, a web-based platform for metabolomics data analysis^105^.

### Data visualization and statistical analysis

Microsoft Office (Microsoft Corporation, Massachusetts, USA) was used for basic graphs. RStudio (Massachusetts, USA) was used to generate transcriptomic heatmaps and volcanic plots with the DESeq2, ggplot2, dplyr, ggrepel, and ComplexHeatmap packages, while multi-omics graphs were created using the mixOmics package. Circos version 0.69-9 was used to visualize transcriptomics and metabolomics data. Statistical bar graphs were generated using GraphPad Prism 10 (GraphPad Software, California, USA). Statistical analyses, including analysis of variance (ANOVA) with a generalized linear model and post-hoc Tukey’s mean comparison test, were performed using Minitab 20 (Minitab, LLC, Pennsylvania, USA). Principal component analysis (PCA) and agglomerative hierarchical clustering (AHC) were performed using XLSTAT Statistical Software 2016 (Addinsoft, Paris, France). Confidence levels were set at 95% and 99%, and data were presented as mean ± SD.

## Author contributions

V.C. conceptualized this study. Y.K. designed, conducted the experiments and analyzed the data under supervision of V.C., P.W., C.W., S.O., A.S. (2). and N.M. D.A designed and conducted the bioinformatic analysis with the help from C.W., P.W. and V.C. A.S. (1). performed the metabolomic experiments. E.M. performed the proteomic experiments. V.C., P.W., N.M., acquired funding and provided project administration. Y.K. and D.A. generated figures with input from all authors. Y.K., D.A. and V.C. wrote the manuscript with input from all authors.

## Acknowledgements

We thank Samantha Murray for her input in the HRC Explorer grant application and Charles Hefer for his help in proteomics experiments. This work was funded by a New Zealand Health Research Council Explorer grant code 22/650 (V.C., N.M. and S.M.), Yili Innovation grant codes 46457 and 46527 (V.C.), KiwiNet Tier 1 grant code 46561 (V.C.) and a Callaghan Innovation PhD Fellowship to Y.K.

## Extended data

**Extended Data Fig. 1.**
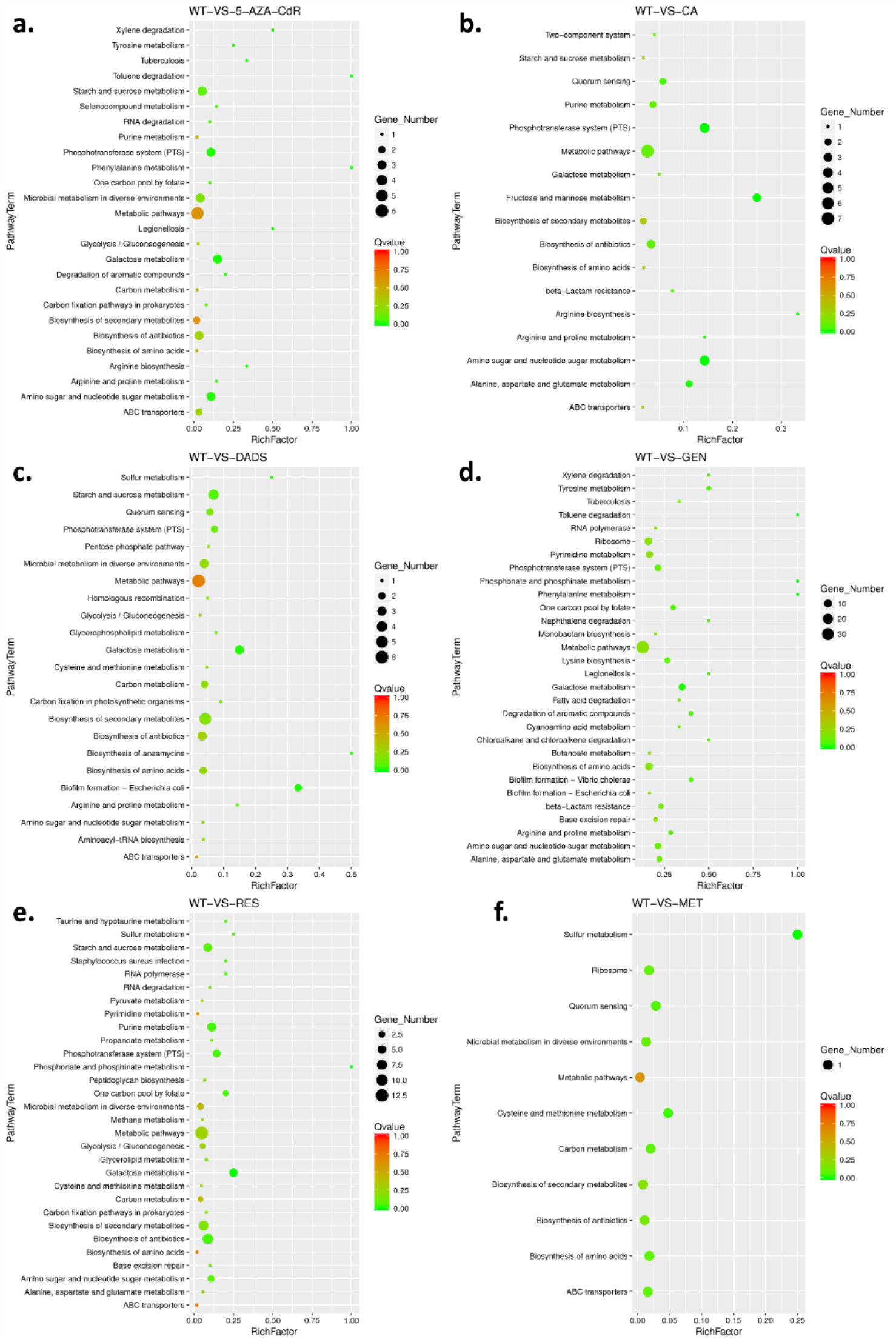
**a-f.** Scatter plots illustrating the KEGG (Kyoto Encyclopedia of Genes and Genomes) pathway enrichment of differentially expressed genes in *Lactobacillus acidophilus*. The X-axis represents the Rich Factor, while the Y-axis lists the specific KEGG pathways. The size of each dot corresponds to the number of differentially expressed genes within the pathway, with larger dots indicating a greater number of genes. The color gradient represents different Q-value ranges, where lower Q-values signify more significant enrichment. Please note that sodium butyrate (SB) was excluded from the KEGG analysis. **Abbreviations:** WT: untreated wild type *L. acidophilus*; 5-AZA-CdR: 5-Aza-2’deoxycytidine; CA: Caffeic Acid; DADS: Diallyl Disulfide; GEN: Genistein; RES: Resveratrol; MET: L-Methionine.

**Extended data Fig. 2.**
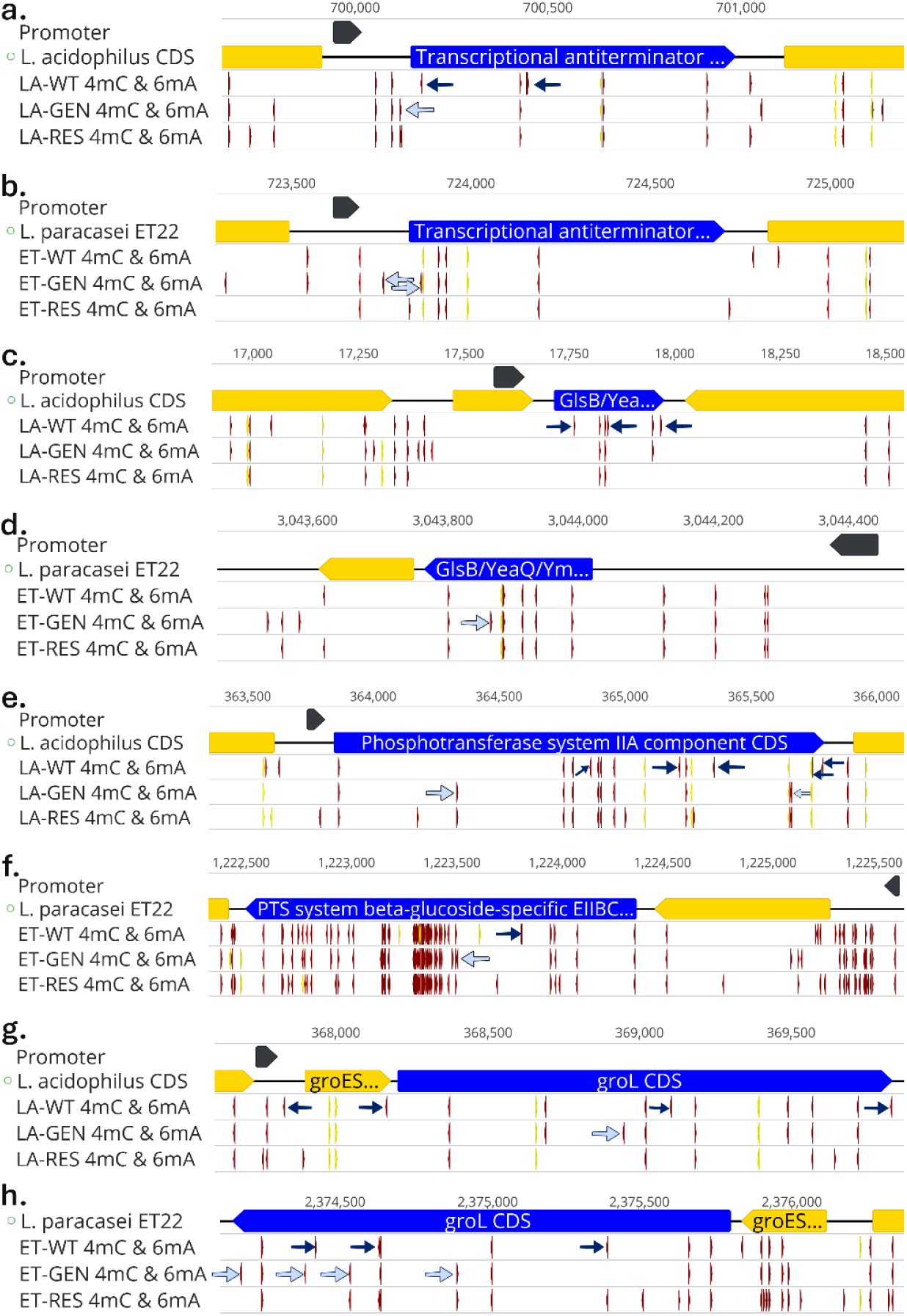
**a-h.** Map of two significantly upregulated genes (a and c, highlighted in blue) and two downregulated genes (e and f, highlighted in blue) in genistein-treated *Lactobacillus acidophilus* (LA-GEN) compared to their homologous genes (b, d, f, and h) in *Lacticaseibacillus paracasei* ET22 (WT). The effect of genistein and resveratrol treatment on the methylation patterns around the promoter region (where predicted) and within the gene body is shown. All panels except panel h include the following: the first track shows the predicted promoter region for the gene, the second track shows the coding sequences in the respective genome (*L. acidophilus* for panels a, c, e, g and *L. paracasei* ET22 for panels b, d, f), the third, fourth, and fifth tracks represent 4mC (in red) and 6mA (in yellow) modification regions in untreated, genistein-treated, and resveratrol-treated *L. acidophilus*, designated LA-WT, LA-GEN, and LA-RES, respectively, and ET-WT, ET-GEN, and ET-RES for *L. paracasei* ET22. The 4mC-modified base(s) in LA-WT but unmodified in LA-GEN is (are) indicated by a dark blue arrow in track three of panels a-g, while the 4mC-modified base(s) in LA-GEN but unmodified in LA-WT is (are) annotated with a light blue arrow in track four of panels a-g, with only one example of each modification type annotated in panel f to maintain clarity. Panel h includes the coding sequences (first track) and 4mC (in red) and 6mA (in yellow) modification regions in untreated, genistein-treated, and resveratrol-treated *L. paracasei* ET22, designated ET-WT, ET-GEN, and ET-RES, respectively (second, third, and fourth tracks).

**Extended data Fig. 3.**
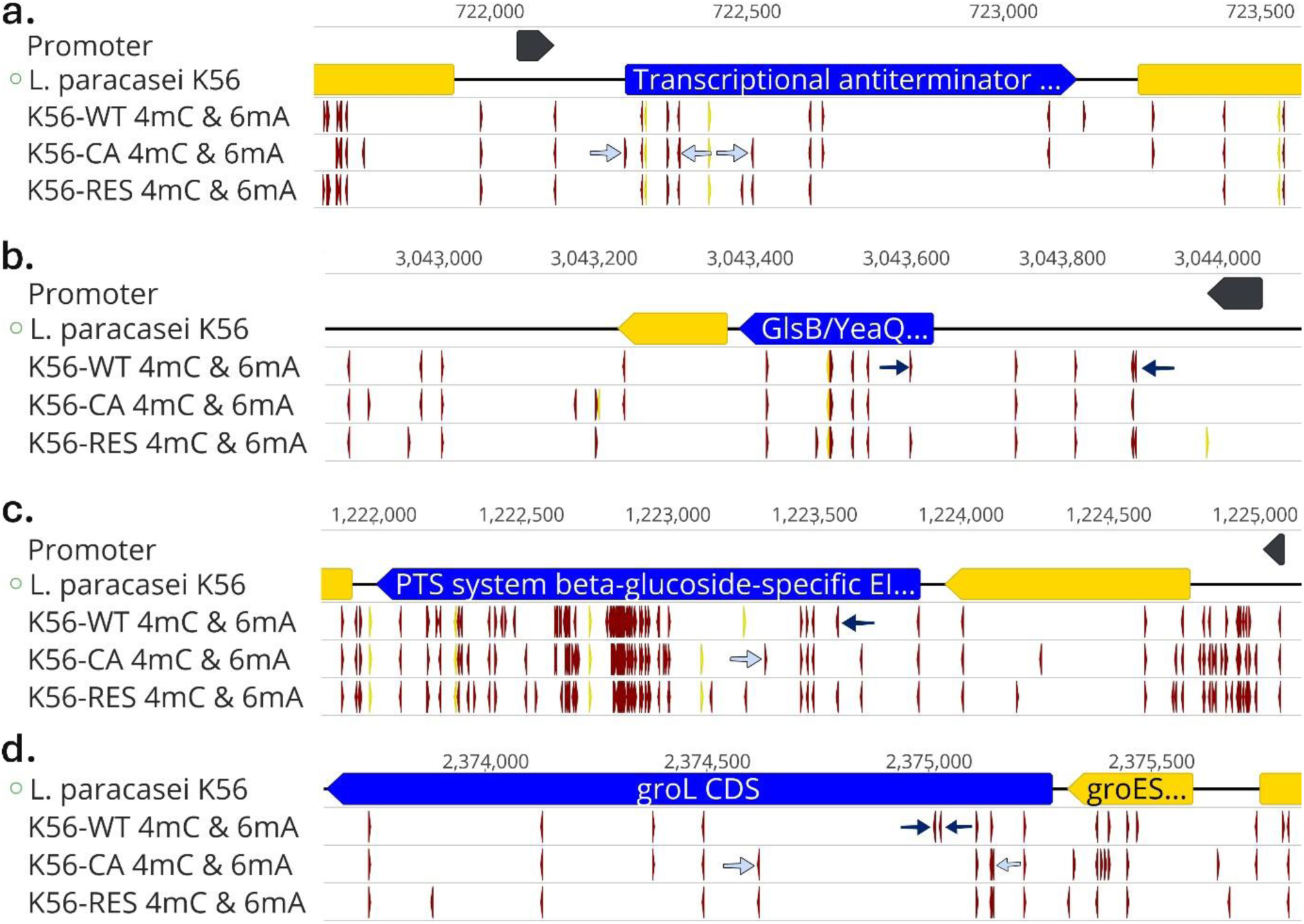
**a-d**. Map of four homologous genes of two significantly upregulated genes (a and b, highlighted in blue) and two downregulated genes (c and d highlighted in blue) in genistein-treated *Lactobacillus acidophilus* in *Lacticaseibacillus paracasei* K56 (K56-WT). The effect of caffeic acid (K56-CA) and resveratrol treatment (K56-RES) on the methylation patterns around the promoter region (where predicted) and within the gene body is shown. Each panel (a-d) includes the following tracks: the first track shows the predicted promoter region for the gene (no promoter predicted for the gene in panel d); the second track shows the coding sequences in *L. paracasei* K56; the third, fourth, and fifth tracks represent 4mC (in red) and 6mA (in yellow) modification regions in untreated, caffeic acid-treated, and resveratrol-treated *L. paracasei,* designated K56-WT, K56-CA, and K56-RES, respectively. The 4mC-modified base(s) in K56-WT but unmodified in K56-CA is(are) indicated by a dark blue arrow in track three of panels a-d. Conversely, the 4mC-modified base(s) in K56-CA but unmodified in K56-WT is(are) annotated with a light blue arrow in track four of panels a-d. Only one example of each modification type is annotated in panel c to maintain clarity.

**Extended Data Fig. 4.**
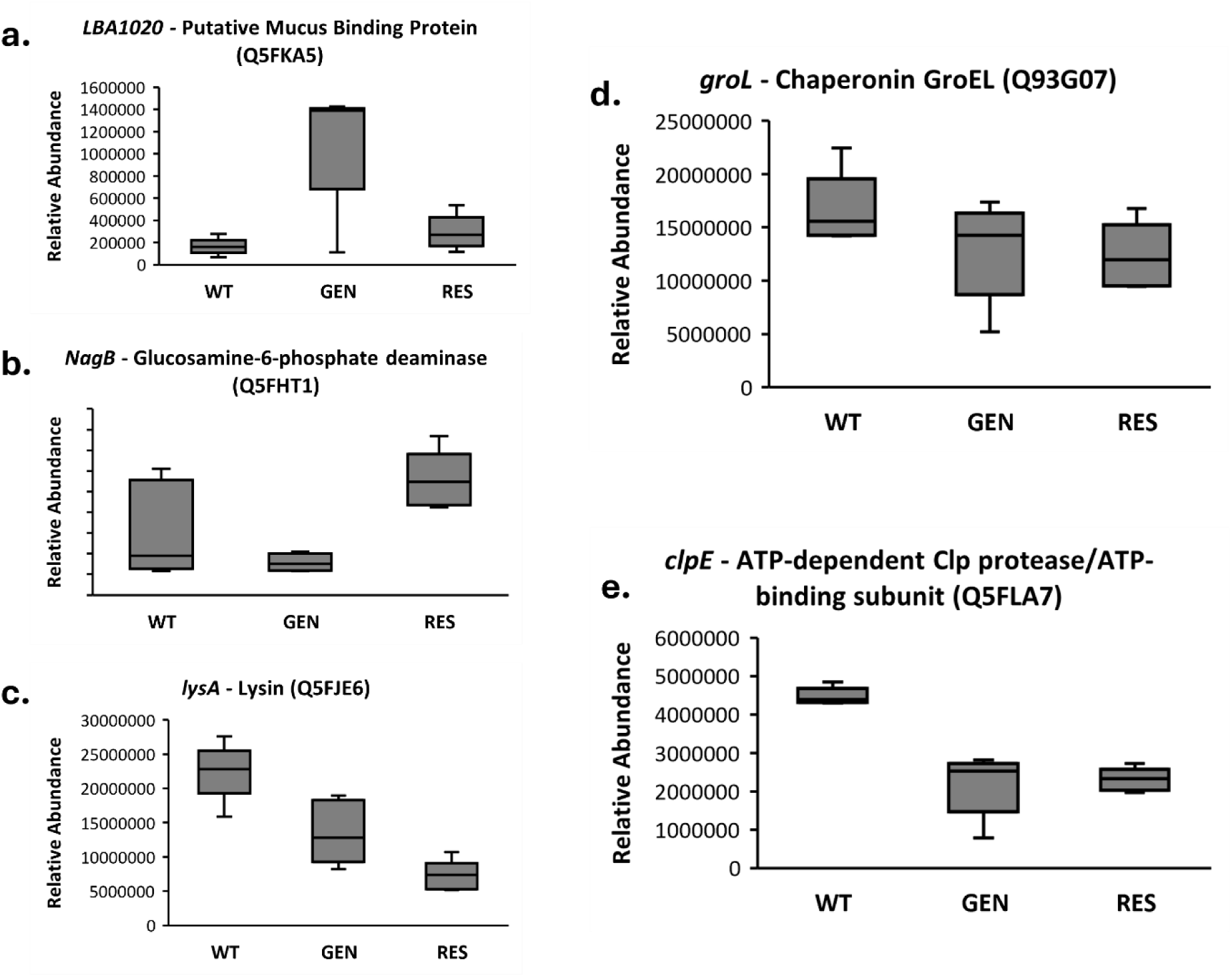
**a-e.** Relative abundance of specific proteins produced by *Lactobacillus acidophilus* under various epigenetic treatments. The selected proteins correspond to genes that were among the top upregulated and downregulated identified by transcriptomic analysis and were also detected in the proteomics data.

**Tukey pairwise mean comparisons (*P* < 0.05): a. *LBA1020* - Putative Mucus Binding Protein (Q5FKA5):** WT (b), GEN (a), RES (b); **b. *NagB* - Glucosamine-6-phosphate deaminase (Q5FHT1):** WT (ab), GEN (b), RES (a); **c. *lysA* - Lysin (Q5FJE6):** WT (a), GEN (b), RES (b); **d. *groL* - Chaperonin GroEL (Q93G07):** WT (a), GEN (a), RES (a); **e**. ***clpE* - ATP-dependent Clp protease/ATP-binding subunit (Q5FLA7):** WT (a), GEN (b), RES (b).

**Extended Data Fig 5.**
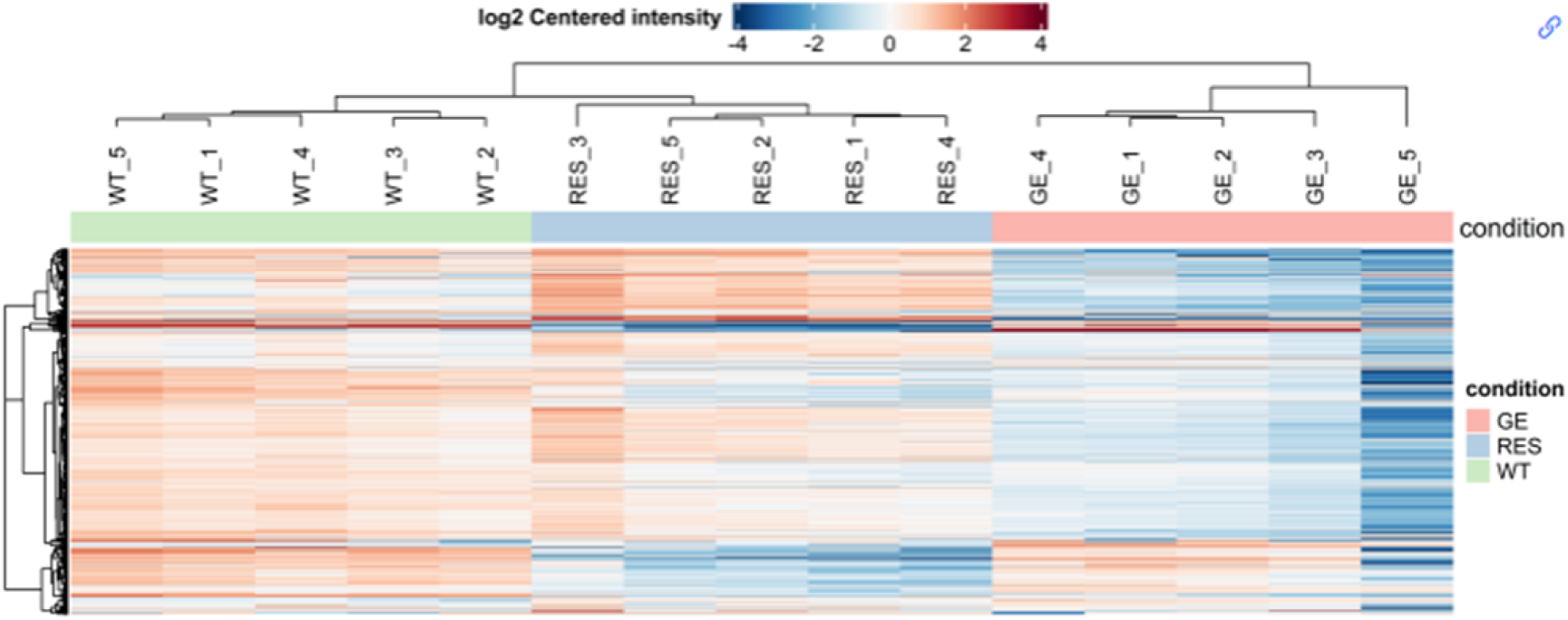
Hierarchical clustering of proteomics results from three different epigenetically treated *Lactobacillus acidophilus* groups: wild type (WT), genistein (GEN), and resveratrol (RES). The heatmap provides a log2-centred measure of expression in each replicate across all conditions, highlighting similarities and differences in expression between experimental contrasts.

**Extended Data Table 1.**
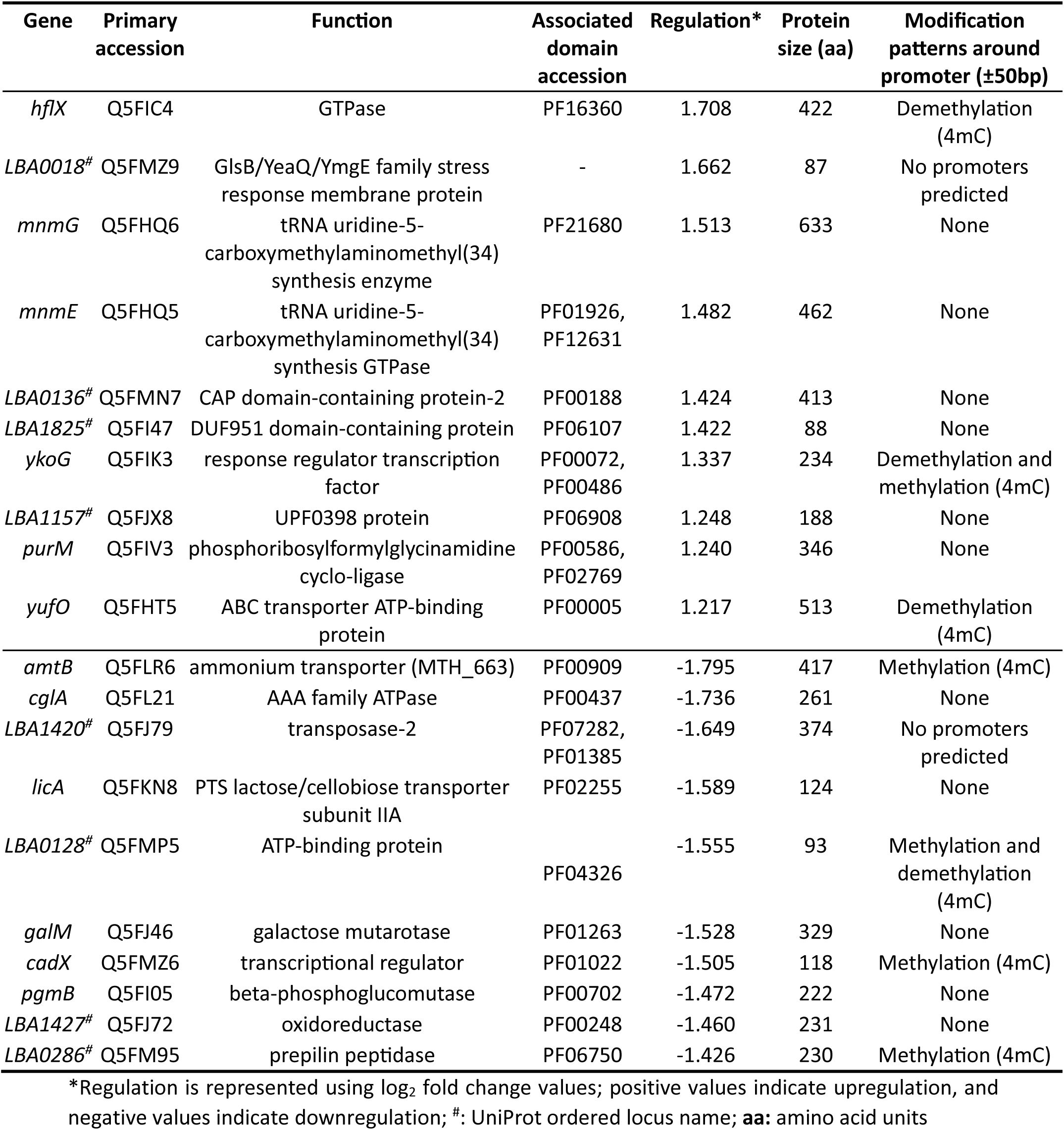
Summary of the top ten significantly upregulated and downregulated genes in resveratrol-treated *L. acidophilus*.

| Gene | Primary accession | Function | Associated domain accession | Regulation* | Protein size (aa) | Modification patterns around promoter ($\pm 50$bp) |
| --- | --- | --- | --- | --- | --- | --- |
| *hflX* | Q5FIC4 | GTPase | PF16360 | 1.708 | 422 | Demethylation (4mC) |
| *LBA0018*<sup>#</sup> | Q5FMZ9 | GlsB/YeaQ/YmgE family stress response membrane protein | - | 1.662 | 87 | No promoters predicted |
| *mnmG* | Q5FHQ6 | tRNA uridine-5-carboxymethylaminomethyl(34) synthesis enzyme | PF21680 | 1.513 | 633 | None |
| *mnmE* | Q5FHQ5 | tRNA uridine-5-carboxymethylaminomethyl(34) synthesis GTPase | PF01926, PF12631 | 1.482 | 462 | None |
| *LBA0136*<sup>#</sup> | Q5FMN7 | CAP domain-containing protein-2 | PF00188 | 1.424 | 413 | None |
| *LBA1825*<sup>#</sup> | Q5FI47 | DUF951 domain-containing protein | PF06107 | 1.422 | 88 | None |
| *ykoG* | Q5FIK3 | response regulator transcription factor | PF00072, PF00486 | 1.337 | 234 | Demethylation and methylation (4mC) |
| *LBA1157*<sup>#</sup> | Q5FJX8 | UPF0398 protein | PF06908 | 1.248 | 188 | None |
| *purM* | Q5FIV3 | phosphoribosylformylglycinamide cyclo-ligase | PF00586, PF02769 | 1.240 | 346 | None |
| *yufO* | Q5FHT5 | ABC transporter ATP-binding protein | PF00005 | 1.217 | 513 | Demethylation (4mC) |
| *amtB* | Q5FLR6 | ammonium transporter (MTH_663) | PF00909 | -1.795 | 417 | Methylation (4mC) |
| *cglA* | Q5FL21 | AAA family ATPase | PF00437 | -1.736 | 261 | None |
| *LBA1420*<sup>#</sup> | Q5FJ79 | transposase-2 | PF07282, PF01385 | -1.649 | 374 | No promoters predicted |
| *licA* | Q5FKN8 | PTS lactose/cellobiose transporter subunit IIA | PF02255 | -1.589 | 124 | None |
| *LBA0128*<sup>#</sup> | Q5FMP5 | ATP-binding protein | PF04326 | -1.555 | 93 | Methylation and demethylation (4mC) |
| *galM* | Q5FJ46 | galactose mutarotase | PF01263 | -1.528 | 329 | None |
| *cadX* | Q5FMZ6 | transcriptional regulator | PF01022 | -1.505 | 118 | Methylation (4mC) |
| *pgmB* | Q5FI05 | beta-phosphoglucomutase | PF00702 | -1.472 | 222 | None |
| *LBA1427*<sup>#</sup> | Q5FJ72 | oxidoreductase | PF00248 | -1.460 | 231 | None |
| *LBA0286*<sup>#</sup> | Q5FM95 | prepilin peptidase | PF06750 | -1.426 | 230 | Methylation (4mC) |

\*Regulation is represented using log<sub>2</sub> fold change values; positive values indicate upregulation, and negative values indicate downregulation; <sup>#</sup>: UniProt ordered locus name; **aa**: amino acid units

**Extended Data Table 2.** Summary of the top ten significantly upregulated and downregulated genes in genistein-treated *L. acidophilus* showing the similarity (grade) and functions of homologous genes found in *Lacticasei. paracasei* ET22.

| Gene | Primary accession | Function (*L. acidophilus*) | Grade of homologous gene *Lacticasei. paracasei* ET22 (%) | Function of homologous gene in *Lacticasei. paracasei* ET22 |
| --- | --- | --- | --- | --- |
| *melR* | Q5FIG1 | Melibiose operon regulatory protein | N/A |  |
| *LBA1020<sup>#</sup>* | Q5FKA5 | Mucin binding | N/A |  |
| *araC* | Q5FME2 | Arabinose operon control; Helix-turn-helix | N/A |  |
| *licT* | Q5FL29 | Transcription antiterminator | 73.5 | transcription antiterminator |
| *LBA1031<sup>#</sup>* | Q5FK95 | Protein kinase | N/A |  |
| *LBA0493<sup>#</sup>* | Q5FLP6 | Aggregation promoting protein | N/A |  |
| *nagB* | Q5FHT1 | Glucosamine-6-phosphate deaminase | 76.6 | Glucosamine-6-phosphate deaminase |
| *LBA0018<sup>#</sup>* | Q5FMZ9 | GlsB/YeaQ/YmgE family stress response membrane protein | 72.8 | Transglycosylase associated protein |
| *lysA* | Q5FJE6 | Lysin | 44.9 | LysM peptidoglycan-binding domain-containing protein |
| *LBA0873<sup>#</sup>* | Q5FKP2 | CsbD-like family protein | N/A |  |
| *LBA0125<sup>#</sup>* | Q5FMP8 | HTH cro/C1-type domain-containing protein | N/A |  |
| *LBA0205<sup>#</sup>* | Q5FMH4 | Heat shock low molecular weight | 68.7 | Hsp20/alpha crystallin family protein |
| *groL* | Q93G07 | Chaperonin GroEL | 87.8 | 60 kDa chaperonin |
| *LBA1606<sup>#</sup>* | Q5FIQ3 | DUF4870 | 43.5 | hypothetical protein Lpp124_14003 |
| *Unnamed* |  | DUF3923 | N/A |  |
| *clpE* | Q5FLA7 | ATPase | 78.7 | ATP-dependent Clp protease |
| *psiD* | Q5FIQ2 | Phosphatidylserine decarboxylase family protein | N/A |  |
| *proC* | Q5FK14 | Pyrroline-5-carboxylate reductase | 45.1 | pyrroline-5-carboxylate reductase |
| *scrA* | G1UB46 | Phosphotransferase system | 74.6 | PTS beta-glucoside transporter subunit |
| *proC* | Q5FK13 | Pyrroline reductase | N/A |  |

**Extended Data Table 3.**
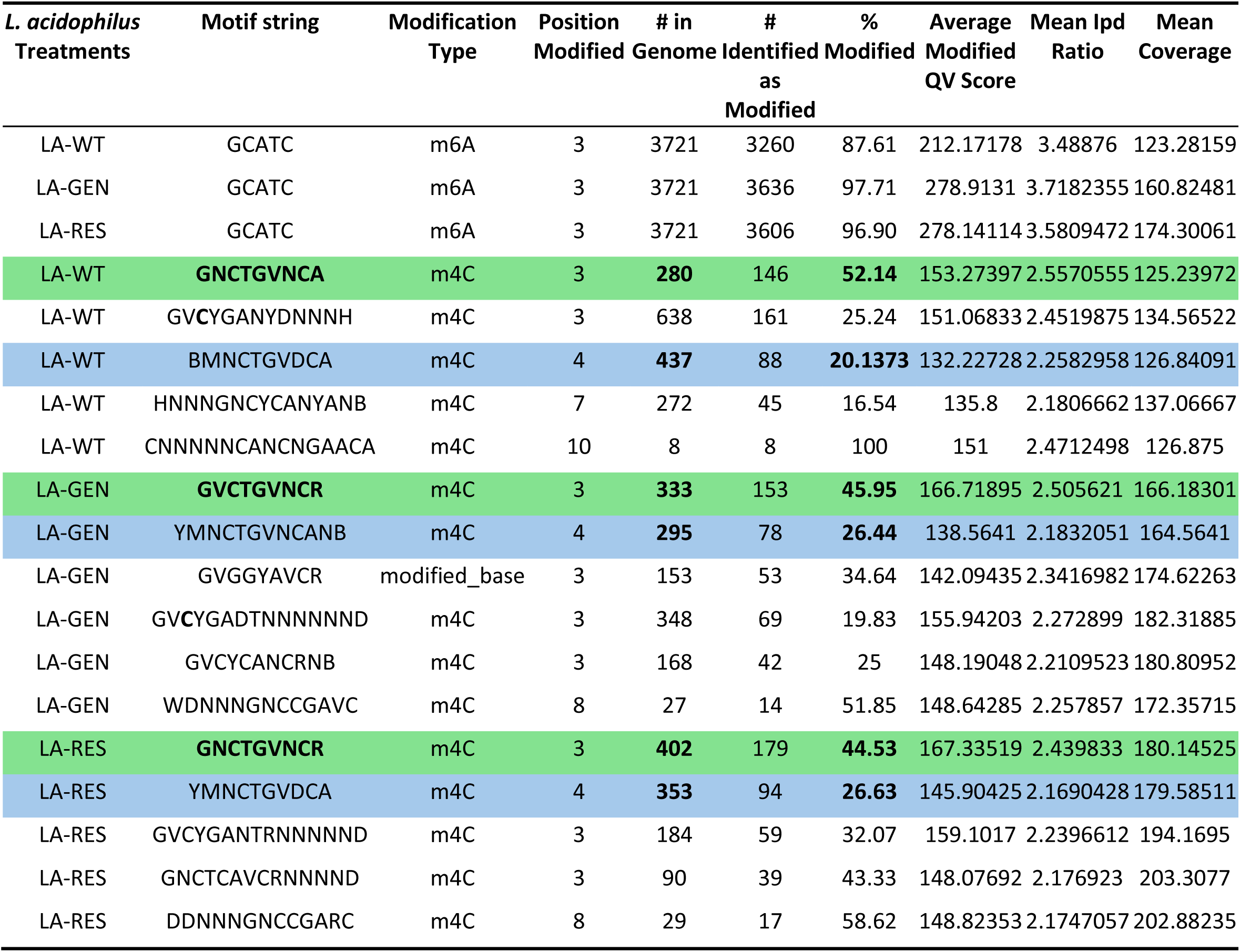
Methylation motifs of untreated *Lactobacillus acidophilus* (LA-WT) vs genistein-treated *L. acidophilus* (LA-GEN) and resveratrol-treated *L. acidophilus* (LA-RES). Two groups of 4mC motifs with similar modification percentages in LA-GEN and LA-RES compared with LA-WT are highlighted in green and blue. Motifs of the same group are highlighted with the same color

## References

1 Hou, K. et al. Microbiota in health and diseases. Signal Transduction and Targeted Therapy 7, 1–28 (2022).

2 Altermann, E. et al. Complete genome sequence of the probiotic lactic acid bacterium *Lactobacillus acidophilus* NCFM. Proceedings of the National Academy of Sciences 102, 3906–3912 (2005).

3 Bull, M., Plummer, S., Marchesi, J. & Mahenthiralingam, E. The life history of *Lactobacillus acidophilus* as a probiotic: a tale of revisionary taxonomy, misidentification and commercial success. FEMS Microbiology Letters 349, 77–87 (2013).

4 Anjum, N., et al. *Lactobacillus acidophilus*: characterization of the species and application in food production. Critical Reviews in Food Science and Nutrition 54, 1241–1251 (2014).

5 Sonnenburg, J. L. & Bäckhed, F. Diet–microbiota interactions as moderators of human metabolism. Nature 535, 56–64 (2016).

6 Sonnenburg, E. D. et al. Diet-induced extinctions in the gut microbiota compound over generations. Nature 529, 212–215 (2016).

7 David, L. A. et al. Diet rapidly and reproducibly alters the human gut microbiome. Nature 505, 559–563 (2014).

8 Kong, Y. et al. Epigenetic Changes in *Saccharomyces cerevisiae* Alters the Aromatic Profile in Alcoholic Fermentation. Applied and Environmental Microbiology 88, e01528–01522 (2022).

9. Ye, P. et al. MethSMRT: an integrative database for DNA N6-methyladenine and N4-methylcytosine generated by single-molecular real-time sequencing. *Nucleic Acids Research*, gkw950 (2016).

10 Luo, G. -Z., Blanco, M. A., Greer, E. L., He, C. & Shi, Y. DNA N 6-methyladenine: a new epigenetic mark in eukaryotes? Nature Reviews Molecular Cell Biology 16, 705–710 (2015).

11 Pisciotta, A. et al. The DNA cytosine methylome revealed two methylation motifs in the upstream regions of genes related to morphological and physiological differentiation in *Streptomyces coelicolor* A (3) 2 M145. Scientific Reports 13, 7038 (2023).

12 Seong, H. J., Han, S.-W. & Sul, W. J. Prokaryotic DNA methylation and its functional roles. Journal of Microbiology 59, 242–248 (2021).

13 Furuta, Y. & Kobayashi, I. in Bacterial integrative mobile genetic elements pp 85–103 (CRC Press, 2022).

14 Fang, J.-L. et al. m4C DNA methylation regulates biosynthesis of daptomycin in *Streptomyces roseosporus* L30. Synthetic and Systems Biotechnology 7, 1013–1023 (2022).

15 Morgan, R. D. et al. Novel m4C modification in type I restriction-modification systems. Nucleic Acids Research 44, 9413–9425 (2016).

16 Kumar, S. et al. N4-cytosine DNA methylation regulates transcription and pathogenesis in *Helicobacter pylori*. Nucleic Acids Research 46, 3429–3445 (2018).

17 Furuta, Y. et al. Methylome diversification through changes in DNA methyltransferase sequence specificity. PLoS Genetics 10, e1004272 (2014).

18 Pan, M. et al. Genomic and epigenetic landscapes drive CRISPR-based genome editing in Bifidobacterium. Proceedings of the National Academy of Sciences 119, e2205068119 (2022).

19 Makarova, K. S. et al. Evolutionary classification of CRISPR–Cas systems: a burst of class 2 and derived variants. Nature Reviews Microbiology 18, 67–83 (2020).

20 Selle, K. & Barrangou, R. Harnessing CRISPR–Cas systems for bacterial genome editing. Trends in Microbiology 23, 225–232 (2015).

21 Wu, Z. et al. Strategies for developing CRISPR-based gene editing methods in bacteria. Small Methods 4, 1900560 (2020).

22 Vento, J. M., Crook, N. & Beisel, C. L. Barriers to genome editing with CRISPR in bacteria. Journal of Industrial Microbiology and Biotechnology 46, 1327–1341 (2019).

23 Arroyo-Olarte, R. D., Bravo Rodriguez, R. & Morales-Ríos, E. Genome editing in bacteria: CRISPR-Cas and beyond. Microorganisms 9, 844 (2021).

24 Kahramanoglou, C., Webster, C. L., El-Robh, M. S., Belyaeva, T. A. & Busby, S. J. Mutational analysis of the *Escherichia coli* melR gene suggests a two-state concerted model to explain transcriptional activation and repression in the melibiose operon. Journal of Bacteriology 188, 3199–3207 (2006).

25 Deutscher, J., Francke, C. & Postma, P. W. How phosphotransferase system-related protein phosphorylation regulates carbohydrate metabolism in bacteria. Microbiology and Molecular Biology Reviews 70, 939–1031 (2006).

26 Nakai, H. et al. The maltodextrin transport system and metabolism in *Lactobacillus acidophilus* NCFM and production of novel α-glucosides through reverse phosphorolysis by maltose phosphorylase. The FEBS Journal 276, 7353–7365 (2009).

27 Rodionova, I. A. et al. The nitrogen regulatory PII protein (GlnB) and N-acetylglucosamine 6-phosphate epimerase (NanE) allosterically activate glucosamine 6-phosphate deaminase (NagB) in *Escherichia coli*. Journal of Bacteriology 200, 10.1128/jb.00691-00617 (2018).

28 Alvarez-Anorve, L. I., Calcagno, M. L. & Plumbridge, J. Why does Escherichia coli grow more slowly on glucosamine than on N-acetylglucosamine? Effects of enzyme levels and allosteric activation of GlcN6P deaminase (NagB) on growth rates. Journal of Bacteriology 187, 2974–2982 (2005).

29 Velasco, A., Leguina, J. & Lazcano, A. Molecular evolution of the lysine biosynthetic pathways. Journal of Molecular Evolution 55, 445–449 (2002).

30 Gaufichon, L., Reisdorf-Cren, M., Rothstein, S. J., Chardon, F. & Suzuki, A. Biological functions of asparagine synthetase in plants. Plant Science 179, 141–153 (2010).

31 Shi, R. et al. Structure-function analysis of *Escherichia coli* MnmG (GidA), a highly conserved tRNA-modifying enzyme. Journal of Bacteriology 191, 7614–7619 (2009).

32 Manickam, N., Joshi, K., Bhatt, M. J. & Farabaugh, P. J. Effects of tRNA modification on translational accuracy depend on intrinsic codon–anticodon strength. Nucleic Acids Research 44, 1871–1881 (2016).

33 Iskandar, C. F., Cailliez-Grimal, C., Borges, F. & Revol-Junelles, A.-M. Review of lactose and galactose metabolism in Lactic Acid Bacteria dedicated to expert genomic annotation. Trends in Food Science and Technology 88, 121–132 (2019).

34 Hall, B. G. The EBG system of *E. coli*: origin and evolution of a novel β-galactosidase for the metabolism of lactose. Genetica 118, 143–156 (2003).

35 Reith, J. & Mayer, C. Peptidoglycan turnover and recycling in Gram-positive bacteria. Applied Microbiology and Biotechnology. 92, 1–11 (2011).

36 Barriere, C. et al. Fructose utilization in *Lactococcus lactis* as a model for low-GC gram-positive bacteria: its regulator, signal, and DNA-binding site. Journal of Bacteriology 187, 3752–3761 (2005).

37 Helanto, M., Aarnikunnas, J., Palva, A., Leisola, M. & Nyyssölä, A. Characterization of genes involved in fructose utilization by *Lactobacillus fermentum*. Archives of Microbiology 186, 51–59 (2006).

38 Bogs, J. & Geider, K. Molecular analysis of sucrose metabolism of *Erwinia amylovora* and influence on bacterial virulence. Journal of Bacteriology 182, 5351–5358 (2000).

39 King, N. D., Hojnacki, D. & O’Brian, M. R. The *Bradyrhizobium japonicum* proline biosynthesis gene proC is essential for symbiosis. Applied and Environmental Microbiology 66, 5469–5471 (2000).

40 Spangler, J. R., Dean, S. N., Leary, D. H. & Walper, S. A. Response of *Lactobacillus plantarum* WCFS1 to the Gram-negative pathogen-associated quorum sensing molecule N-3-oxododecanoyl homoserine lactone. Frontiers in Microbiology 10, 715 (2019).

41 Jordan, S., Hutchings, M. I. & Mascher, T. Cell envelope stress response in Gram-positive bacteria. FEMS Microbiology Reviews 32, 107–146 (2008).

42 Zellmeier, S., Schumann, W. & Wiegert, T. Involvement of Clp protease activity in modulating the *Bacillus subtilis* σW stress response. Molecular Microbiology 61, 1569–1582 (2006).

43 Damjanovic, M., Kharat, A. S., Eberhardt, A., Tomasz, A. & Vollmer, W. The essential tacF gene is responsible for the choline-dependent growth phenotype of *Streptococcus pneumoniae*. Journal of Bacteriology 189, 7105–7111 (2007).

44 Haijema, B. J., Hahn, J., Haynes, J. & Dubnau, D. A ComGA-dependent checkpoint limits growth during the escape from competence. Molecular Microbiology 40, 52–64 (2001).

45 Lee, W.-H., Shin, S.-Y., Kim, M.-D., Han, N. S. & Seo, J.-H. Modulation of guanosine nucleotides biosynthetic pathways enhanced GDP-L-fucose production in recombinant *Escherichia coli*. Applied Microbiology and Biotechnology 93, 2327–2334 (2012).

46 Stoyanov, J. V., Hobman, J. L. & Brown, N. L. CueR (YbbI) of *Escherichia coli* is a MerR family regulator controlling expression of the copper exporter CopA. Molecular Microbiology 39, 502–512 (2001).

47 Le Coq, D., Lindner, C., Krüger, S., Steinmetz, M. & Stülke, J. New beta-glucoside (bgl) genes in *Bacillus subtilis*: the bglP gene product has both transport and regulatory functions similar to those of BglF, its *Escherichia coli* homolog. Journal of Bacteriology 177, 1527–1535 (1995).

48 Brunner, K. et al. Inhibitors of the cysteine synthase CysM with antibacterial potency against dormant *Mycobacterium tuberculosis*. Journal of Medical Chemistry 59, 6848–6859 (2016).

49 Agren, D., Schnell, R., Oehlmann, W., Singh, M. & Schneider, G. Cysteine synthase (CysM) of *Mycobacterium tuberculosis* is an O-phosphoserine sulfhydrylase: evidence for an alternative cysteine biosynthesis pathway in mycobacteria. Journal of Biological Chemistry 283, 31567–31574 (2008).

50 Spoering, A. L. & Gilmore, M. S. Quorum sensing and DNA release in bacterial biofilms. Current Opinions in Microbiology 9, 133–137 (2006).

51 Wang, R. et al. Gut microbiota regulates acute myeloid leukaemia via alteration of intestinal barrier function mediated by butyrate. Nature Communications 13, 2522 (2022).

52 Sealy, L. & Chalkley, R. The effect of sodium butyrate on histone modification. Cell 14, 115–121 (1978).

53 Silva, A. et al. L-pyroglutamic acid inhibits energy production and lipid synthesis in cerebral cortex of young rats in vitro. Neurochemical Research 26, 1277–1283 (2001).

54 Zielke, H. R., Zielke, C. L. & Ozand, P. T. Glutamine: a major energy source for cultured mammalian cells. in Federation Proceedings Vol. 43 121–125 (1984).

55 Randall, K., Lever, M., Peddie, B. A. & Chambers, S. T. Natural and synthetic betaines counter the effects of high NaCl and urea concentrations. Biochimica et Biophysica Acta (BBA)-General Subjects 1291, 189–194 (1996).

56 Chen, W. et al. Absence of the biliverdin reductase-a gene is associated with increased endogenous oxidative stress. Free Radical Biology and Medicine 115, 156–165 (2018).

57 Deguchi, K. et al. Reduction of cerebral infarction in rats by biliverdin associated with amelioration of oxidative stress. Brain Research 1188, 1–8 (2008).

58 Gallego-Lobillo, P., Ferreira-Lazarte, A., Hernández-Hernández, O., Montilla, A. & Villamiel, M. Evaluation of the impact of a rat small intestinal extract on the digestion of four different functional fibers. Food and Function 11, 4081–4089 (2020).

59 Zhou, Y. et al. Efficiency analysis and mechanism Insight of that whole-cell biocatalytic production of melibiose from raffinose with *Saccharomyces cerevisiae*. Applied Biochemistry and Biotechnology. 181, 407–423 (2017).

60 Gabai, G., Mongillo, P., Giaretta, E. & Marinelli, L. Do dehydroepiandrosterone (DHEA) and its sulfate (DHEAS) play a role in the stress response in domestic animals? Frontiers in Veterinary Science 7, 588835 (2020).

61 Omura, Y. Beneficial effects & side effects of DHEA: True anti-aging & age-promoting effects, as well as anti-cancer & cancer-promoting effects of DHEA evaluated from the effects on the normal & cancer cell telomeres & other parameters. Acupuncture and Electrotherapy Research 30, 219–261 (2005).

62 Yoshida, S. et al. Dehydroepiandrosterone sulfate is inversely associated with sex-dependent diverse carotid atherosclerosis regardless of endothelial function. Atherosclerosis 212, 310–315 (2010).

63 Tsimogiannis, D. & Oreopoulou, V. Free radical scavenging and antioxidant activity of 5, 7, 3′, 4′-hydroxy-substituted flavonoids. Innovative Food Science and Emerging Technologies 5, 523–528 (2004).

64 Dixon, R. A. & Ferreira, D. Genistein. Phytochemistry 60, 205–211 (2002).

65 Sekowska, A. et al. Bacterial variations on the methionine salvage pathway. BMC Microbiology 4, 1–17 (2004).

66 Sanderson, S. M., Gao, X., Dai, Z. & Locasale, J. W. Methionine metabolism in health and cancer: a nexus of diet and precision medicine. Nature Reviews Cancer 19, 625–637 (2019).

67 Mu, W., Yu, S., Zhu, L., Zhang, T. & Jiang, B. Recent research on 3-phenyllactic acid, a broad-spectrum antimicrobial compound. Applied Microbiology and Biotechnology 95, 1155–1163 (2012).

68 Rodríguez, N., Salgado, J. M., Cortés, S. & Domínguez, J. M. Antimicrobial activity of D-3-phenyllactic acid produced by fed-batch process against *Salmonella enterica*. Food Control 25, 274–284 (2012).

69 Srikhanta, Y. N. et al. Phasevarions mediate random switching of gene expression in pathogenic Neisseria. PLoS Pathogens 5, e1000400 (2009).

70 Ishikawa, K., Fukuda, E. & Kobayashi, I. Conflicts targeting epigenetic systems and their resolution by cell death: novel concepts for methyl-specific and other restriction systems. DNA Research 17, 325–342 (2010).

71 Furuta, Y. & Kobayashi, I. Mobility of DNA sequence recognition domains in DNA methyltransferases suggests epigenetics-driven adaptive evolution. Mobile Genetic Elements 2, 292–296 (2012).

72 Schiffer, C. J., Grätz, C., Pfaffl, M. W., Vogel, R. F. & Ehrmann, M. A. Characterization of the *Staphylococcus xylosus* methylome reveals a new variant of type I restriction modification system in staphylococci. Frontiers in Microbiology 14, 946189 (2023).

73 Roberts, R. J., Vincze, T., Posfai, J. & Macelis, D. REBASE—a database for DNA restriction and modification: enzymes, genes and genomes. Nucleic Acids Research 43, D298–D299 (2015).

74 Kelleher, P., Bottacini, F., Mahony, J., Kilcawley, K. N. & van Sinderen, D. Comparative and functional genomics of the *Lactococcus lactis* taxon; insights into evolution and niche adaptation. BMC Genomics 18, 1–20 (2017).

75 Kumari, M., Swarnkar, M. K., Kumar, S., Singh, A. K. & Gupta, M. Complete genome sequence of potential probiotic *Lactobacillus sp*. HFC8, isolated from human gut using PacBio SMRT sequencing. Genome Announcements 3, 10.1128/genomea.01337-01315 (2015).

76 Sturino, J. M., Rajendran, M. & Altermann, E. Draft genome sequence of *Lactobacillus animalis* 381-IL-28. Genome Announcements 2, 10.1128/genomea.00478-00414 (2014).

77 Gulati, P., Singh, A., Goel, M. & Saha, S. The extremophile *Picrophilus torridus* carries a DNA adenine methylase M. PtoI that is part of a Type I restriction-modification system. Frontiers in Microbiology 14, 1126750 (2023).

78 Furuta, Y., Abe, K. & Kobayashi, I. Genome comparison and context analysis reveals putative mobile forms of restriction–modification systems and related rearrangements. Nucleic Acids Research 38, 2428–2443 (2010).

79 Khan, F. et al. A putative mobile genetic element carrying a novel type IIF restriction-modification system (PluTI). Nucleic Acids Research 38, 3019–3030 (2010).

80 Furuta, Y. & Kobayashi, I. Movement of DNA sequence recognition domains between non-orthologous proteins. Nucleic Acids Research 40, 9218–9232 (2012).

81. 81 Wei, X., et al. Vaginal microbiomes show ethnic evolutionary dynamics and positive selection of *Lactobacillus* adhesins driven by a long-term niche-specific process. Cell Reports 43 (2024).

82 Segal, E. et al. Module networks: identifying regulatory modules and their condition-specific regulators from gene expression data. Nature Genetics 34, 166–176 (2003).

83 Xu, W. et al. Enzymatic production of melibiose from raffinose by the levansucrase from *Leuconostoc mesenteroides* B-512 FMC. Journal of Agricultural and Food Chemistry 65, 3910–3918 (2017).

84 İspirli, H., Kaya, Y. & Dertli, E. Bifidogenic effect and in vitro immunomodulatory roles of melibiose-derived oligosaccharides produced by the acceptor reaction of glucansucrase E81. Process Biochemistry 91, 126–131 (2020).

85 Hullar, M. A. & Fu, B. C. Diet, the gut microbiome, and epigenetics. The Cancer Journal 20, 170–175 (2014).

86 Wang, X. et al. Antioxidant and anti-inflammatory effects of 6, 3’, 4-and 7, 3, 4-Trihydroxyflavone on 2D and 3D RAW264. 7 Models. Antioxidants 12, 204 (2023).

87 Matsuda, H., Morikawa, T., Ando, S., Toguchida, I. & Yoshikawa, M. Structural requirements of flavonoids for nitric oxide production inhibitory activity and mechanism of action. Bioorganic and Medicinal Chemistry 11, 1995–2000 (2003).

88 Alexandre, L. S. et al. Flavonoids, cytotoxic, and antimalarial activities of *Dipteryx lacunifera*. Revista Brasileira de Farmacognosia 30, 544–550 (2020).

89 Suzuki, M. M. & Bird, A. DNA methylation landscapes: provocative insights from epigenomics. Nature Reviews Genetics 9, 465–476 (2008).

90 Lee, W. C. et al. The complete methylome of Helicobacter pylori UM032. BMC Genomics 16, 1–9 (2015).

91 Zhao, J. et al. Roles of adenine methylation in the physiology of *Lacticaseibacillus paracasei*. Nature Communications 14, 2635 (2023).

92 Nurk, S. et al. HiCanu: accurate assembly of segmental duplications, satellites, and allelic variants from high-fidelity long reads. Genome research 30, 1291–1305 (2020).

93 Walker, B. J. et al. Pilon: an integrated tool for comprehensive microbial variant detection and genome assembly improvement. PloS One 9, e112963 (2014).

94 Schwengers, O. et al. Bakta: rapid and standardized annotation of bacterial genomes via alignment-free sequence identification. Microbial Genomics 7, 000685 (2021).

95 Aziz, R. K. et al. The RAST Server: Rapid Annotations using Subsystems Technology. BMC Genomics 9, 75, doi:10.1186/1471-2164-9-75 (2008).

96 Arkin, A. P. et al. KBase: the United States department of energy systems biology knowledgebase. Nature Biotechnology 36, 566–569 (2018).

97 UniProt Consortium. UniProt: the universal protein knowledgebase in 2023. Nucleic Acids Research 51, D523–D531 (2023).

98 Rice, P., Longden, I. & Bleasby, A. EMBOSS: the European molecular biology open software suite. Trends in Genetics 16, 276–277 (2000).

99 Crooks, G. E., Hon, G., Chandonia, J.-M. & Brenner, S. E. WebLogo: a sequence logo generator. Genome Research 14, 1188–1190 (2004).

100 de Jong, A., Pietersma, H., Cordes, M., Kuipers, O. P. & Kok, J. PePPER: a webserver for prediction of prokaryote promoter elements and regulons. BMC Genomics 13, 299, doi:10.1186/1471-2164-13-299 (2012).

101 Benjamini, Y. & Hochberg, Y. Controlling the false discovery rate: a practical and powerful approach to multiple testing. Journal of the Royal Statistical Society: series B (Methodological) 57, 289–300 (1995).

102 Rohart, F., Gautier, B., Singh, A. & Lê Cao, K.-A. mixOmics: An R package for ‘omics feature selection and multiple data integration. PLoS Computational Biology 13, e1005752, doi:10.1371/journal.pcbi.1005752 (2017).

103 Ritchie, M. E. et al. limma powers differential expression analyses for RNA-sequencing and microarray studies. Nucleic Acids Research 43, e47–e47 (2015).

104 Tsugawa, H. et al. MS-DIAL: data-independent MS/MS deconvolution for comprehensive metabolome analysis. Nature Methods 12, 523–526 (2015).

105 Pang, Z. et al. MetaboAnalyst 5.0: narrowing the gap between raw spectra and functional insights. Nucleic Acids Research 49, W388–W396 (2021).

